# HSD17B13 Couples Hepatocyte Lipid Metabolism to Stellate Cell Activation via TGFb-1 Signaling

**DOI:** 10.1101/2025.10.14.682191

**Authors:** Naideline Raymond, Lawrence Lifshitz, Kyounghee Min, Caroline A. Lewis, Giorgis Isaac, Batuhan Yenilmez, Michael P. Czech

## Abstract

Metabolic dysfunction-associated steatohepatitis (MASH) is a progressive liver disease driven by hepatocellular lipid overload, immune activation, and hepatic stellate cell (HSC)-driven fibrogenesis. Human genetic studies reveal that loss-of-function (LoF) variants in 17 beta-hydroxysteroid dehydrogenase 13 (*HSD17B13*) confer robust protection against advanced fibrosis and cirrhosis, establishing *HSD17B13* as a critical genetic modifier of MASH severity. Yet the mechanisms linking HSD17B13 activity to fibrogenic progression remain poorly understood. Here, we show that both wild-type and catalytically deficient HSD17B13 (mHSD) localize to lipid droplets (LDs) in cultured human hepatocytes, but only catalytically active HSD17B13 enhances hepatocellular lipid accumulation and markedly upregulates the lipogenic transcriptional regulator carbohydrate-responsive element-binding protein (ChREBP). This HSD17B13-driven lipogenic axis elicits potent paracrine activation of LX2 stellate cells, both in hepatocyte-HSC co-culture and in response to hepatocyte-conditioned medium (CM). Screening of candidate signaling mediators revealed that transforming growth factor beta-1 (TGFb-1) is uniquely and strongly upregulated by active HSD17B13, with minimal induction by mHSD. Remarkably, siRNA-mediated knockdown of *TGFB1* or neutralization of active TGFb-1 protein abolishes CM-induced LX2 activation and collagen synthesis. Collectively, these findings identify HSD17B13 as a dual metabolic and profibrotic effector that drives TGFb-1-dependent HSC activation, thereby linking hepatocellular lipid dysregulation to fibrogenic progression and providing a mechanistic framework for understanding how HSD17B13 contributes to MASH pathogenesis.

## Introduction

Metabolic dysfunction–associated steatotic liver disease (MASLD) has emerged as the most prevalent chronic liver disorder worldwide, affecting nearly 30% of adults [1–4]. MASLD spans a clinical spectrum from simple steatosis to its progressive inflammatory form, metabolic dysfunction–associated steatohepatitis (MASH), which can advance to severe fibrosis, cirrhosis, and hepatocellular carcinoma (HCC) [5–8]. The rising incidence of MASLD parallels the global epidemics of obesity, type 2 diabetes (T2D), and related cardiometabolic disorders, posing a growing global health challenge [3, 4, 9–11].

Therapeutic development has accelerated in recent years, as two pharmacological agents have gained FDA approval for the treatment of noncirrhotic MASH with moderate to advanced fibrosis. Resmetirom (Rezdiffra), a liver-targeted thyroid hormone receptor-β (THR-β) agonist, was approved in 2024 [12–15] and semaglutide (Wegovy) received accelerated approval in 2025 for histologic resolution of MASH [16, 17]. Although both agents demonstrate histologic efficacy, their antifibrotic effects remain modest. No current therapy fully halts or reverses advanced liver fibrosis, underscoring a critical unmet need for interventions that directly and robustly target fibrogenic pathways [14, 18–21]. Meanwhile, multiple therapeutic classes are under active clinical evaluation, including THR-β agonists, glucagon-like peptide-1 (GLP-1)/ glucose-dependent insulinotropic polypeptide (GIP) receptor agonists, fibroblast growth factor 21 (FGF21) analogs, farnesoid X receptor (FXR) and pan-peroxisome proliferator– activated receptor (pan-PPAR) agonists, each targeting distinct pathogenic nodes in MASH such as dysregulated lipid metabolism, chronic inflammation, and fibrogenesis [13, 22–25]. In this context, our laboratory is actively developing preclinical strategies that jointly target steatotic and fibrogenic pathways in experimental models of MASH [26].

Despite these advances, a deeper mechanistic understanding of hepatic lipid partitioning and the crosstalk among hepatocytes, immune cells, and hepatic stellate cells (HSCs) in MASH is essential to identify key pathogenic regulators and guide the development of durable disease-modifying therapies. It is well established that excessive accumulation of toxic lipid species in hepatocytes triggers oxidative stress and inflammatory signaling, which together drive hepatocellular injury and initiate fibrogenesis through activation of HSCs—the principal source of extracellular matrix (ECM) in the liver [27–35]. However, the molecular pathways that couple hepatocellular steatosis to fibrogenic activation remain incompletely defined.

Potential clues to these mechanisms have emerged from human genetic studies showing that loss-of-function (LoF) variants in 17β-hydroxysteroid dehydrogenase 13 (*HSD17B13*) confer protection against lobular inflammation, advanced fibrosis, and cirrhosis [36–43]. HSD17B13 is selectively enriched in hepatocytes and localizes to lipid droplets (LDs) which are dynamic organelles that coordinate lipid storage, metabolism, and signaling [36, 44–48]. When dysregulated, LD-associated proteins can promote lipid accumulation, mitochondrial dysfunction, and inflammatory signaling that characterize MASLD and MASH [49–53]. These observations have spurred intense interest in *HSD17B13* as a therapeutic target, with RNA interference-based approaches already in early-phase clinical trials [54–57] and small-molecule inhibitors of HSD17B13 enzymatic activity showing efficacy in preclinical models [47, 58, 59]. Nonetheless, how LD-localized HSD17B13 shapes hepatocellular metabolism to promote profibrotic signaling remains unresolved.

Extensive alternative splicing of *HSD17B13* yields at least nine isoforms, among which isoform A (IsoA) represents the full-length, catalytically active form [46, 60]. HSD17B13 is an NAD(P)H/NAD(P)+-dependent oxidoreductase that catalyzes the reversible interconversion of retinoids, steroids, and other bioactive lipids [46, 48, 60–62]. Despite these biochemical insights, functional studies in mice have produced results that diverge from human genetic data. *Hsd17b13* knockout mice fail to recapitulate the hepatoprotective phenotype observed in human LoF carriers, likely reflecting species-specific differences in enzymatic function and substrate preference [63–68]. Notably, murine Hsd17b13 appears to lack retinol dehydrogenase (RDH) activity, an enzymatic function retained in the human ortholog and implicated in the regulation of retinoic acid (RA) signaling. Additional evidence suggests that the closely related paralog *Hsd17b11* may compensate for *Hsd17b13* loss in mice [64]. By contrast, *HSD17B13* is more abundantly expressed than *HSD17B11* in human liver, reducing the likelihood of functional redundancy [62, 64].

Mechanistically, HSD17B13 has been implicated in both lipolytic suppression and lipogenic activation. HSD17B13 has been shown to inhibit adipose triglyceride lipase (ATGL)-dependent lipolysis by directly associating with ATGL and preventing its activation by the co-factor α-β hydrolase domain-containing protein 5 (ABHD5/CGI-58) on the LD surface [68] and has also been linked to enhanced activation of sterol regulatory element-binding protein 1c (SREBP-1c), a master regulator of hepatic lipogenesis [69]. While these studies suggest that HSD17B13 influences lipid handling, through both enzymatic activity and structural interactions at the LD surface, the downstream signaling consequences of its expression remain unknown. A deeper understanding of how the catalytic and scaffolding functions of HSD17B13 contribute to fibrogenic progression in human models is essential to connect genetic protection with mechanistic insight.

This study was designed to define the role of HSD17B13 as a modifier of fibrosis risk and to determine whether its catalytic activity is required for hepatic lipid accumulation and profibrotic signaling. Here, we show that HSD17B13 catalytic activity drives both lipid accumulation and a hepatocyte HSD17B13→ transforming growth factor beta 1 (TGFb-1) signaling axis that promotes paracrine activation of human HSCs. Given that TGFb-1 is a master regulator of fibrogenic responses [70–77], these findings establish a mechanistic link between hepatocellular lipid dysregulation and fibrogenesis, highlighting HSD17B13 as a dual metabolic and profibrotic driver in MASLD/MASH.

## Materials and Methods

### Plasmid DNA Electroporation

Expression vectors encoding human *HSD17B13* with an N-terminal HA-tag under the control of a CMV promoter were obtained from VectorBuilder. The catalytically inactive (S172A) mutant was generated by substituting serine at position 172 with alanine to abrogate *HSD17B13* enzymatic activity [46]. A non-coding open reading frame (negative ORF) vector served as a transfection control. Plasmid DNA (3–5 μg) was diluted in Resuspension Buffer R and electroporated into Huh7 or HepG2 cells using the Neon™ Transfection System (Thermo Fisher Scientific, Cat. #MPK10096), per manufacturer’s instructions. Electroporation was performed under optimized parameters (1350 V, 30 ms, single pulse) [78]. Cells were immediately transferred into pre-warmed complete growth medium, consisting of Dulbecco’s Modified Eagle Medium (DMEM; Gibco, Cat# 11965-092) supplemented with 10% fetal bovine serum (FBS; Gibco), and 1% penicillin-streptomycin (100 U/mL penicillin, 100 µg/mL streptomycin). Cells were then incubated overnight at 37°C in 5% CO₂ prior to downstream analyses.

### Huh7 Cell Culture and Studies

Huh7 human hepatoma cells (Glow Biologics, Cat# GBTC-099H) were maintained in complete growth medium. Cells were cultured at 37°C and passaged at 80–90% confluency using 0.25% trypsin-EDTA (Gibco). Experiments were conducted using cells between passages 2 and 12. For experiments, Huh7 cells were seeded in 6-, 12-, or 24-well standard plates at densities appropriate for the downstream assay and allowed to adhere overnight. For lipid loading experiments, cells were treated with vehicle, 100 µM of bovine serum albumin (BSA)-complexed with palmitate (PA) and oleate (OA) (1PA2OA), and/or 5 µM retinol (ROL) for 48–72 h, as indicated.

### LX2 Human Hepatic Stellate Cell Culture and Studies

LX2 human hepatic stellate cells (HSCs; Millipore Sigma, Cat. #SCC064) were cultured in complete growth medium. Cells were serum-starved overnight before stimulation. For direct co-culture experiments, LX2 cells were plated in 6- or 12-well Transwell plates (Thermo Fisher, Cat. #140642 and 140654), or in Pair-N-Share™ Tandem Co-Culture Wells (Bulldog Bio, Cat. #ARPS001, ARPS002, ARPSAP1) assembled per the manufacturer’s instructions. Pair-N-Share wells were fitted with a 1.2 µm filters (Bulldog Bio, Cat. #ARPSF003) to physically separate LX2 and Huh7 hepatocytes while allowing paracrine signaling. For hepatocyte-conditioned media (CM) experiments, LX2 cells were plated in standard culture plates and treated with hepatocyte-CM for 48 h before harvest. As a positive control for HSC activation, serum-starved LX2 cells were stimulated with 5 ng/mL recombinant human TGFb-1 (R&D Systems, Cat. #240-B/CF) for 48 h.

### Primary Human Hepatocyte (PHH) Culture and Studies

Cryopreserved PHHs (BioIVT) were thawed and handled per the supplier’s protocol. Cells were resuspended in InVitroGRO™ CP medium (BioIVT, Cat. #Z990003) supplemented with TORPEDO antibiotic mix (BioIVT, Cat. #Z99008) and 10% FBS. Cells were plated onto collagen I-coated wells or coverslips at a density and kept in fresh CP medium. After 4–6 h of attachment, medium was replaced with InVitroGRO™ HI maintenance medium (BioIVT, Cat. #Z990012) containing 0.1% FBS. Medium was refreshed every 24–48 h. Cells were treated with vehicle, 100 µM 1PA2OA, and/or 5 µM ROL for up to 72 h, followed by downstream analyses.

### Human HepG2 Hepatoma Culture and Studies

HepG2 human hepatoma cells (ATCC, HB-8065) were cultured in RPMI-1640 medium (Gibco, Cat# 11875-093) supplemented with 10% FBS. For lipid-accumulation assays, cells were seeded onto glass coverslips in 24-well plates and incubated overnight. The following day, cells were treated as indicated with vehicle, 100 µM 1PA2OA, and/or 5 µM ROL for 48—72 h before Oil Red O staining.

### ORO Staining and Brightfield Microscopy

Cells were washed twice with Dulbecco’s phosphate-buffered saline (DPBS) and fixed in 4% paraformaldehyde for 15 min at room temperature. Cells were then rinsed and incubated in 60% isopropanol for 5 min. Filtered ORO working solution (0.3% w/v in isopropanol and diluted 3:2 in water) was applied to the cells for 15 min at room temperature. Stained cells were rinsed and mounted with aqueous medium. Images were acquired under identical exposure settings on a brightfield microscope (40× objective). Lipid-positive areas were quantified using ImageJ (NIH), with thresholding applied to the red channel.

### Triglyceride (TG) Quantification

Intracellular TGs were measured using the Cayman Kit (Cat# 10010303) per manufacturer’s instructions. Cells were washed and lipids were extracted with 3:2 hexane:isopropanol solution for 30 min. The organic solvent was evaporated, and lipids were resuspended. Samples or TG standards (10µL) were loaded in duplicate in a 96-well plate and incubated with the enzymatic reaction mixture for 1 h. The absorbance was measured at 540 nm on safire2 microplate reader. TG concentrations were calculated from a standard curve and expressed as fold-change relative to mock controls.

### RNA Isolation, cDNA Synthesis and Quantitative Real-Time PCR (RT-qPCR)

Total RNA was isolated using TRIzol™ Reagent (Invitrogen, Cat# 15596026) following the manufacturer’s protocol. RNA pellets were washed with 75% ethanol, air-dried, and resuspended in RNase-free water. RNA concentration and purity were determined by NanoDrop spectrophotometry. cDNA synthesis was performed with the iScript™ cDNA Synthesis Kit (Bio-Rad, Cat# 1708891) using 1 µg RNA. RT-qPCR was carried out using iTaq™ Universal SYBR® Green Supermix (Bio-Rad, Cat# 1725121) on a CFX96 Touch™ system. Reactions were run in technical triplicate, and gene expression was analyzed using the 2^−ΔΔCt method normalized to *GAPDH*.

### SDS-PAGE and Immunoblotting of Cultured Cells

Cells were lysed in ice-cold radioimmunoprecipitation assay (RIPA) buffer (50 mM Tris-HCl, pH 7.4; 150 mM NaCl; 1% NP-40; 0.5% sodium deoxycholate; 0.1% sodium dodecyl sulfate (SDS)) supplemented with protease/phosphatase inhibitors (Sigma-Aldrich). Protein concentrations were measured using the Pierce BCA Protein Assay. Equal amounts of protein were denatured at 95°C for 5-10 min and resolved on 4-16% gradient polyacrylamide gels (Bio-Rad). Proteins were transferred to nitrocellulose membranes and blocked in Tris-buffered saline with 0.1% Tween-20 (TBST) with 5% non-fat dry milk or 5% BSA for 1 h. Membranes were then incubated overnight at 4°C with primary antibodies (1:200—1:2000). After washing, membranes were incubated with HRP-conjugated secondary antibodies (1:5000) for 1 h at room temperature. Bands were visualized by enhanced chemiluminescence (PerkinElmer/Revvity) using a Bio-Rad ChemiDox XRS+ system. Band intensities were quantified in Fiji/ImageJ and normalized to housekeeping proteins (Histone H3, HSP90, or Vinculin).

### TGFb-1 Quantification by ELISA (Thermo Fisher)

Total human TGFb-1 levels were measured using the Human TGFb-1 ELISA Kit (Thermo Fisher Scientific, Cat# BMS249-4). To quantify total TGFb-1, prediluted samples were acid-activated 1 N HCl for 1 h and neutralized with 1 N NaOH. Standards, controls, and samples were loaded in duplicate to 96-well microplates pre-coated with a monoclonal anti–TGFb-1 antibody. Samples were then washed and incubated with biotinylated detection antibody for 1 h, followed by streptavidin–horseradish peroxidase (HRP) for 1 h. Colorimetric detection was performed with the tetramethyl-benzidine (TMB) substrate for 25-30 min and stopped with 1M phosphoric acid. The absorbance was measured at 450 nm. Concentrations of TGFb-1 were calculated from a standard curve and normalized to total protein.

### Cell-Based Retinol Dehydrogenase Assay for HSD17B13 Activity

Huh7 cells were transfected with *HSD17B13* or control plasmids and cultured overnight. The next day, cells were incubated in serum-free medium containing 1 µM all-trans retinol for 8 h. Medium and cells were collected separately, and retinoids were extracted using ice-cold 50:50 (v/v) acetonitrile/water solution containing deuterium-labeled internal standards, followed by methyl tert-butyl ether (MTBE) extraction. The upper organic phase was collected, dried, resuspended in 1:3 water: methanol, and analyzed by LC-MS analysis (UMass Metabolomics Core).

### Cell-Based Lipolysis Assay

Glycerol release was measured using the Cayman Glycerol Cell-Based Assay Kit (Cat# 10011725). Collected culture supernatant or glycerol standards (25 µL) were incubated in duplicate with reconstituted assay reagent. Following incubation, absorbance was read at 540 nm, and concentrations were calculated from a glycerol standard curve.

### Immunofluorescence (IF) Assay

Cells were seeded on coverslips and allowed to adhere overnight. Cells were washed and fixed with 4% paraformaldehyde for 15 min. Fixed cells were permeabilized with 0.1% saponin (Sigma) and blocked for 1 h in filtered normal goat blocking solution containing 2% goat serum, 1% BSA, 0.1% triton X-100, and 0.05% tween 20. Cells were incubated overnight at 4°C with primary antibodies (1:100– 1:500) in a humidified chamber. Cells were then washed and incubated for 1 h with Alexa Fluor 647-conjugated secondary antibodies (1:500). Neutral lipids were stained with LipidTOX (1:500) for 30-60 min, and nuclei were counterstained with DAPI (1 µg/mL). Coverslips were mounted with ProLong Gold Antifade (Thermo Fisher, cat P36931). Images were acquired on a Leica SP8 confocal microscope with appropriate filters.

### In vitro silencing of TGFb-1 in hepatocytes

Huh7 cells were transfected with mock, IsoA, or mHSD constructs and allowed to recover overnight. Cells were then transfected with 1 µM Accell™ siRNA targeting *TGFB1* (Dharmacon, Cat# E-003929-00) or non-targeting control (NTC) siRNA for 24 h. After 24 h, cells were washed and treated with a 1PA2OA and 5 µM ROL for 48 h. Cells were harvested for RNA isolation and *TGFB1* expression analysis. Hepatocyte-CM were collected and applied to serum-starved LX2 cells for 48 h prior to downstream assays. For TGFb-1 neutralization experiment, hepatocyte-CM was incubated overnight with anti-TGFb-1 antibody (1 μg/mL) or isotype control before application to LX2 cells.

### Quantification and statistical analysis

Statistical analyses were performed using GraphPad Prism 9. Comparisons between two groups were analyzed by two-tailed unpaired Student’s *t*-test. Multiple-group comparisons involved one-way or two-way ANOVA when appropriate. Data are presented as mean ± SEM unless otherwise stated. Statistical significance was defined as *p* < 0.05 and annotated as: *p* < 0.05 (*) (^#^), *p* < 0.01 (**) (^##^), *p* < 0.001 (***) (^###^), and *p* < 0.0001 (****) (^####^). Outliers were identified using Prism’s built-in outlier test and excluded only with clear technical justification. Experiments were repeated at least twice with ≥3 biological replicates unless otherwise noted. Graphs display individual data points to illustrate replicate variability.

## RESULTS

### HSD17B13 Associates with LDs and Promotes Lipid Accumulation in Hepatocytes

To define the function of *HSD17B13* in hepatocytes, we ectopically expressed the *HSD17B13* wild-type isoform (IsoA) in hepatoma cell lines lacking endogenous expression. Immunoblotting confirmed robust IsoA expression at levels comparable to endogenous HSD17B13 protein expression observed in PHHs (Fig. 1A). Next, we examined the effect of HSD17B13 on LD dynamics and steatosis. Huh7 cells transfected with IsoA were cultured for 48 h in media supplemented with a palmitate (PA) and oleate (OA) mixture (1:2 ratio, final concentration 100 μM; hereafter referred to as 1PA2OA) and 5 μM retinol (ROL), a known HSD17B13 substrate [62]. Hepatocytes maintained in 10% FBS medium served as controls. LDs were visualized using LipidTOX™ staining and quantified with ImageJ. IsoA protein (red) localized to the LD surface (Fig. 1B), consistent with prior reports that HSD17B13 is an LD-associated protein [36, 44–48].

**Figure 1:**
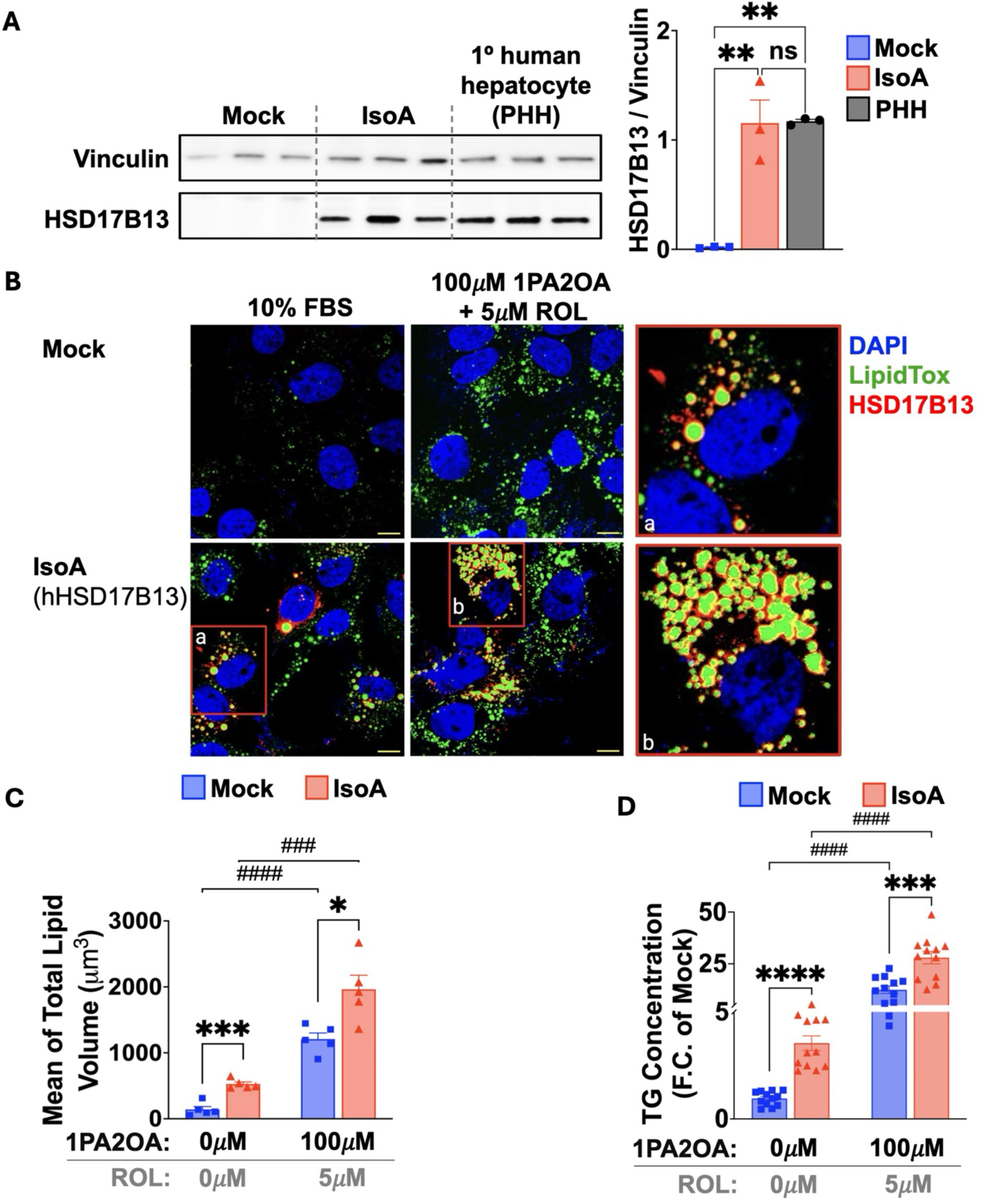
LD-associated HSD17B13 promotes hepatocellular lipid accumulation. **(A)** Immunoblot analysis confirmed robust expression of wild-type *HSD17B13* (IsoA) in Huh7 cells following plasmid transfection, at levels comparable to endogenous HSD17B13 expression in PHHs. **(B)** Huh7 cells transfected with IsoA were cultured for 48 h in 10% FBS medium or in medium supplemented with 100 μM 1PA2OA and 5 μM ROL. Immunostaining showed IsoA localization to the LD surface (red), with LDs stained by LipidTox Green (green) and nuclei counterstained with DAPI (blue). Scale bars, 20 μm. **(C)** Confocal images were quantified using ImageJ to measure LD volume, expressed as normalized LipidTox Green fluorescence intensity. IsoA overexpression significantly increased lipid accumulation under both basal and lipid-loaded conditions compared with mock-transfected cells. **(D)** Biochemical TG quantification corroborated imaging data, showing elevated TG levels in IsoA-expressing Huh7 cells under both basal and lipid-loading conditions. Data are presented as mean ± SEM. (A—D) *n* ≥ 3 biologically independent samples. Statistical significance was determined by ordinary one-way ANOVA followed by Sidack’s multiple-comparison test and by unpaired two-tailed Student’s *t*-test for individual treatment group comparisons. ***ns*** *= P>0.05;* **\*** *= P≤0.05;* **\*\*** *= P≤0.01;* **\*\*\*** *= P≤0.001;* **\*\*\*\*** *= P≤0.0001*.

IsoA expression significantly increased intracellular lipid accumulation in Huh7 cells under basal conditions, and this effect was further amplified in response to 1PA2OA and ROL supplementation (Fig. 1C). Triglyceride (TG) quantification corroborated these findings as we detected higher TG levels in IsoA-expressing Huh7 cells under both basal and supplemented conditions (Fig. 1D). Moreover, these results were consistent across human hepatocyte cell systems. IsoA overexpression in HepG2 cells enhanced lipid accumulation under both basal and supplemented conditions, as assessed by Oil Red O (ORO) staining (Fig. S1A, B). In PHHs, lipid accumulation observed under basal condition was minimal but increased significantly upon treatment with 100 μM 1PA2OA (Fig. S1C). Dose-response measurements in PHHs showed that increasing concentrations of 1PA2OA not only promoted lipid accumulation but also induced *HSD17B13* mRNA expression, whereas protein levels remained largely unchanged at concentrations below 500 μM (Fig. S1D, E). Collectively, these findings largely suggest that LD-associated HSD17B13 promotes hepatic lipid accumulation in human hepatoma cell lines.

### HSD17B13 Enzymatic Function Drives Maximal Hepatocellular TG Accumulation

A recent study identified key functional domains within *HSD17B13* and pinpointed serine 172 (Ser172) as a critical residue for its catalytic activity [46]. To dissect the contribution of enzymatic activity to HSD17B13 function, we introduced the same serine-to-alanine substitution at residue 172 (S172A) to generate a catalytically inactive mutant (mHSD). Western blot analysis confirmed comparable expression of wild-type IsoA and mHSD in Huh7 cells (Fig. 2A).

**Figure 2:**
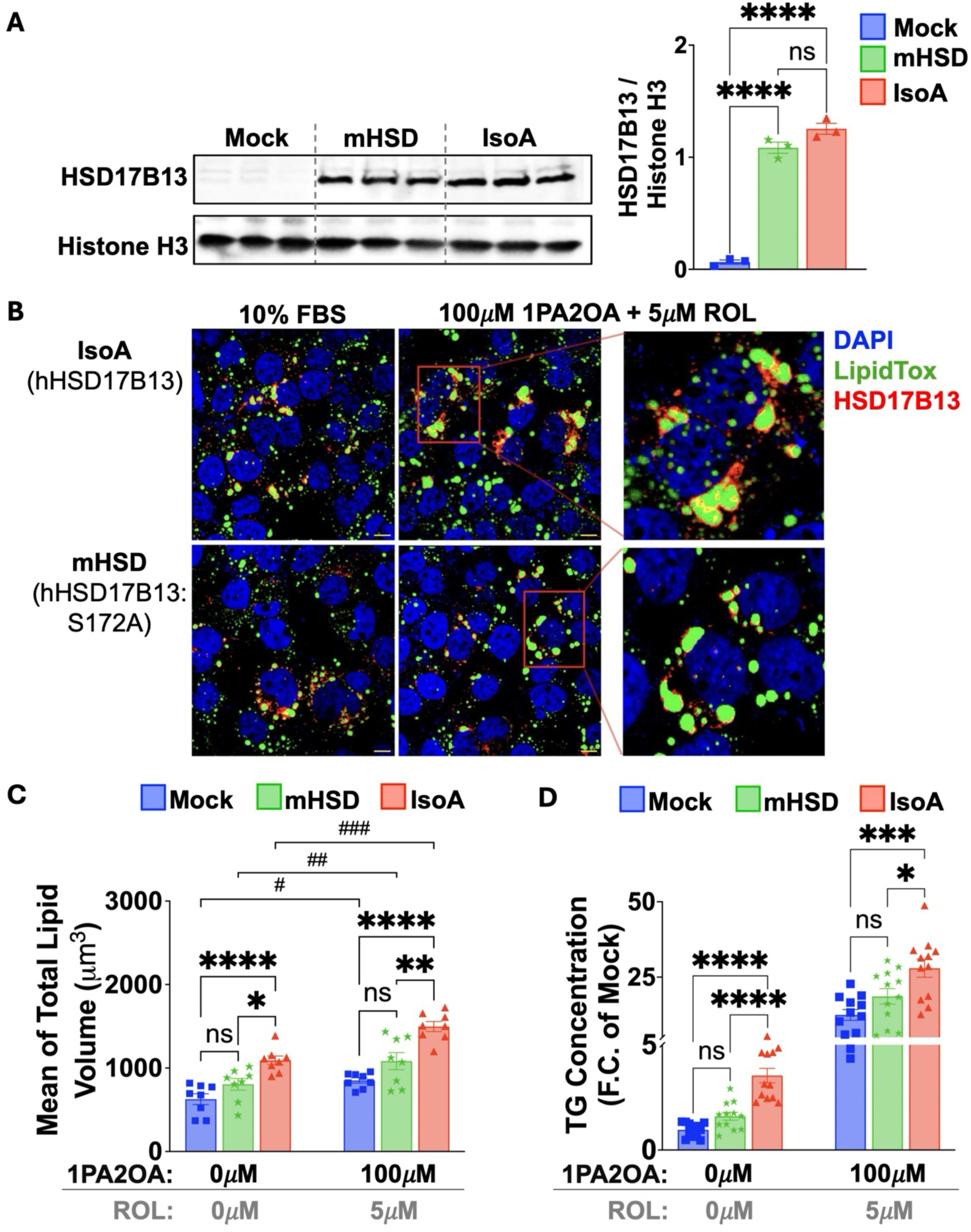
The enzymatic activity of HSD17B13 drives maximal hepatocellular TG accumulation. **(A)** Schematic representation of the HSD17B13 protein highlighting Ser172 as a critical residue for catalytic activity. Western blot analysis confirmed comparable expression of wild-type HSD17B13 (IsoA) and the catalytically inactive mutant HSD17B13-S172A (mHSD) in Huh7 cells following plasmid transfection. **(B)** Huh7 cells expressing IsoA or mHSD were cultured for 48 h in media containing 10% FBS or supplemented with 100 μM 1PA2OA and 5 μM ROL. Immunostaining showed that mHSD localized to LDs (red) similarly to IsoA, confirming that the catalytically inactive mutant retained LD targeting function. LDs were stained by LipidTox Green (green) and nuclei were counterstained with DAPI (blue). Scale bars, 20 μm. **(C)** Quantification of confocal images revealed that IsoA overexpression markedly increased LD volume and TG accumulation under both basal and lipid-loading conditions. In contrast, mHSD expression failed to promote TG accumulation, and remained comparable to mock-transfected controls even in 1PA2OA + ROL–supplemented medium. **(D)** Biochemical TG quantification paralleled imaging data, showing that IsoA expression increased intracellular TG content under both basal and lipid-loaded condition. mHSD expression failed to promote lipid accumulation, with lipid volume and TG levels remaining comparable with mock-transfected controls. Data presented as mean ± SEM. (A—D) *n* ≥ 3 biologically independent samples. Statistical significance between individual treatment group was determined by one-way ANOVA followed by Turkey’s multiple-comparison test. ***ns*** *= P>0.05;* **\*** *= P≤0.05;* **\*\*** *= P≤0.01;* **\*\*\*** *= P≤0.001;* **\*\*\*\*** *= P≤0.0001*.

Because HSD17B13 is thought to possess retinol dehydrogenase (RDH) activity that catalyzes the oxidation of retinol to retinaldehyde (retinal), we evaluated enzymatic function using a cell-based retinol conversion assay. IsoA overexpression increased retinal production, whereas mHSD failed to catalyze retinol oxidation (Fig. S2A). Consistently, IsoA, but not mHSD, induced expression of the retinoic acid (RA)-responsive gene retinaldehyde dehydrogenase (*ALDH1A1*), which encodes the enzyme that converts retinal to RA (Fig. S2B). These findings confirmed that the S172A mutation compromised HSD17B13 catalytic activity without affecting protein stability.

To determine whether catalytic activity is required for neutral lipid enrichment, IsoA- or mHSD- expressing cells were cultured in 10% FBS basal medium with or without 1PA2OA and ROL supplementation for 48 h. Neutral LDs were visualized by LipidTOX™ staining. The S172A mutation did not affect mHSD localization to LDs (Fig. 2C). Yet, whereas IsoA overexpression markedly increased intracellular TG content under both basal and supplemented conditions, mHSD overexpression failed to promote neutral lipid deposition (Fig. 2D). TG levels in mHSD-expressing cells remained comparable to mock controls even in 1PA2OA plus ROL-enriched medium (Fig. 2D). Together, these results define a direct requirement for HSD17B13 enzymatic function in promoting hepatocellular lipid accumulation.

### HSD17B13-Driven Activation of Lipogenic Pathways Requires Catalytic Activity

To further investigate how HSD17B13 promotes neutral lipid accumulation, we examined its effect on lipid synthesis and degradation, the two major processes that regulate LD turnover (Fig. S3A). We first assessed lipolysis by quantifying glycerol release from IsoA- and mHSD-expressing Huh7 cells. Under lipid overload conditions, glycerol release was significantly reduced in both IsoA- and mHSD-transfected cells, indicating that HSD17B13-mediated suppression of lipolysis does not require catalytic activity. At baseline, however, IsoA expression led to a more pronounced impairment in lipolytic activity (Fig. S3B).

We next examined molecular markers of the de novo lipogenesis (DNL) pathway, including key transcription factors and target enzymes. Specifically, we assessed activation of sterol regulatory element-binding protein-1 (SREBP-1) and expression of carbohydrate response element-binding protein (ChREBP) by immunoblotting. No significant differences in SREBP-1 activation were observed among mock-, mHSD-, and IsoA-transfected Huh7 cells, calculated as the ratio of mature [(m)SREBP-1] to its full-length precursor (Fig. S3C). In contrast, ChREBP protein levels were markedly increased in IsoA-transfected cells, with the lowest protein expression observed in mock controls (Fig. S3C). In line with these findings, IsoA overexpression significantly upregulated mRNA expression of DNL-associated enzymes, including fatty acid synthase (*FASN*), stearoyl-CoA desaturase 1 (*SCD1*), acetyl-CoA carboxylase alpha (*ACACA*) and ATP citrate lyase (*ACLY*), whereas mHSD did not affect expression of these genes (Fig. S3D). Taken together, these results suggest that LD–associated HSD17B13 suppresses lipid mobilization partly independent of its catalytic activity but requires catalytic function to promote transcriptional upregulation of lipogenic enzymes, thereby coordinately inhibiting lipolysis and stimulating lipogenic pathways to favor intracellular lipid accumulation in hepatocytes.

### Hepatocyte HSD17B13 Catalytic Function Triggers Paracrine HSC activation and Fibrogenic Signaling

How hepatocyte lipid accumulation in obesity signals to HSCs to drive their activation and subsequent fibrogenesis remains an incompletely understood aspect of MASH pathogenesis, partly due to the heterogeneous manifestations of MASH. To address this gap, we established a human hepatocyte– HSC co-culture model using a horizontal transwell system (Fig. 3A). Huh7 hepatocytes transfected with mock, mHSD, or IsoA constructs were cultured overnight in basal medium. Transfected Huh7 cells were then transitioned to co-culture with immortalized human HSCs (LX2) for 48 h in media supplemented with 100 μM 1PA2OA and 5 μM ROL, with 10% FBS medium serving as basal control. In parallel, LX2 monocultures were maintained under identical conditions to control for direct effects of 1PA2OA and ROL supplementation.

**Figure 3:**
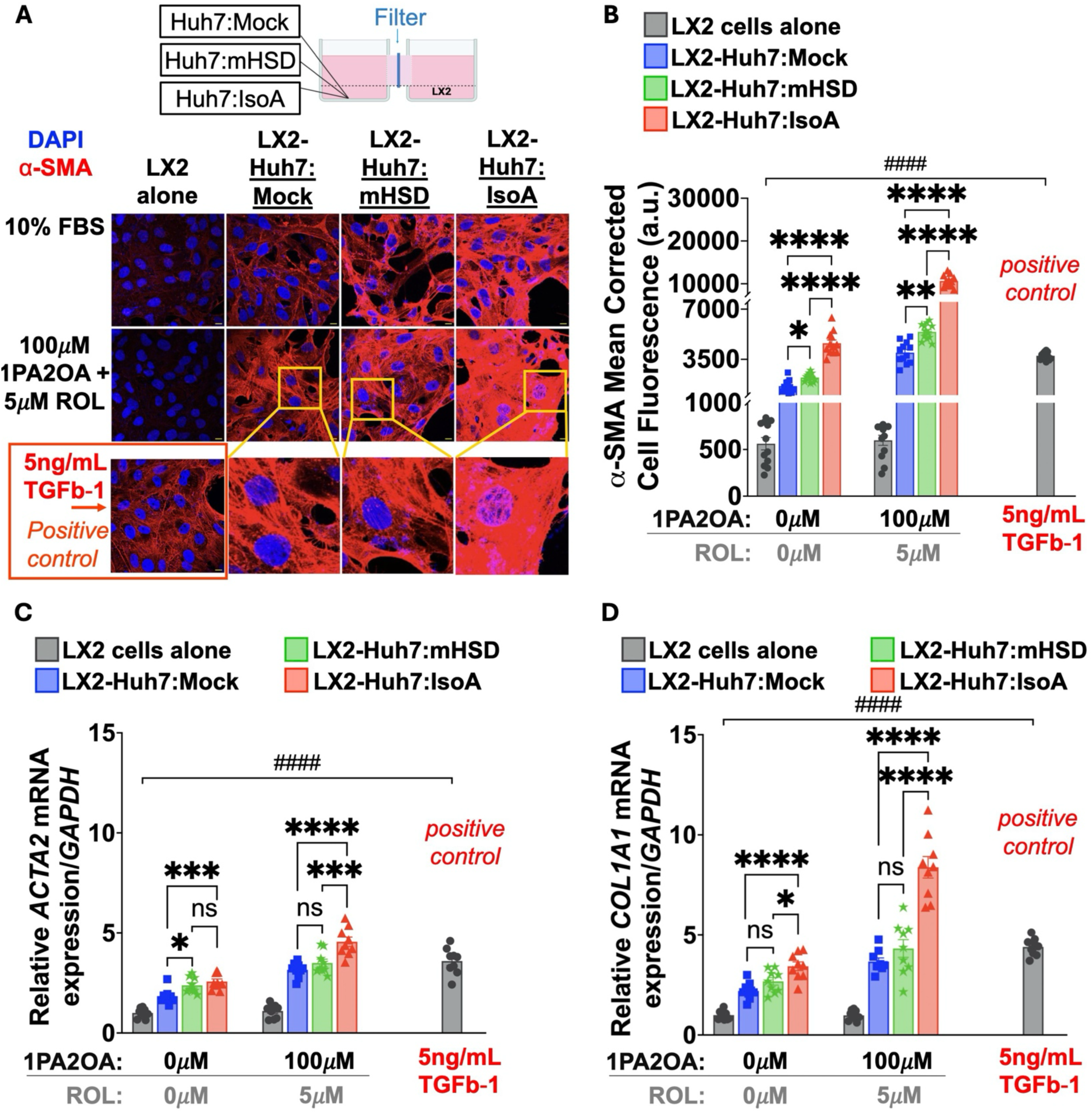
Hepatocyte HSD17B13 catalytic function triggers paracrine HSC activation and fibrogenic signaling in *in vitro* co-culture. **(A)** Schematic of the human hepatocyte–HSC co-culture model using a horizontal transwell system. Huh7 hepatocytes transfected with mock, IsoA, or mHSD constructs were maintained in isolated cultures and then co-cultured with LX2 cells for 48 h. LX2 activation was assessed by immunofluorescence staining for α-smooth muscle actin (α-SMA; red) with DAPI nuclear counterstain (blue). Scale bars, 10 μm. **(B)** Quantification of confocal images reveal that lipid supplementation to LX2 control monocultures did not α-SMA induction, whereas the positive control, recombinant TGFb-1 (5 ng/mL) robustly activated LX2 cells and increased α-SMA induction. Co-culture with Huh7 hepatocytes markedly induced α-SMA expression that rivaled the response elicited by TGFb-1, with IsoA-transfected hepatocytes triggering the strongest α-SMA induction in LX2 cells under both basal and lipid-supplemented conditions. **(C–D)** Quantitative RT-PCR analysis of fibrogenic gene expression in LX2 cells isolated from co-cultures. **(C)** *ACTA2* mRNA (encoding α-SMA) was strongly upregulated in LX2 cells co-cultured with IsoA-transfected hepatocytes under lipid-rich conditions, whereas mock- and mHSD-transfected hepatocytes failed to induce robust induction of *ACTA2*. **(D)** *COL1A1* expression (encoding the α1 chain of type I collagen) was selectively upregulated in LX2 cells co-cultured with IsoA-transfected hepatocytes under both basal and lipid-loaded conditions. Data presented as mean ± SEM. (B-D) *n* ≥ 3 independent samples. Statistical significance between individual treatment group was determined by one-way ANOVA followed by Tukey’s multiple-comparison test. ***ns*** *= P>0.05;* **\*** *= P≤0.05;* **\*\*** *= P≤0.01;* **\*\*\*** *= P≤0.001;* **\*\*\*\*** *= P≤0.0001*.

Because induction of alpha smooth muscle actin (α-SMA, encoded by *ACTA2*) is a hallmark of HSC activation, we assessed LX2 activation by immunofluorescence using a fluorogenic anti-α-SMA antibody (red) and DAPI nuclear counterstain (blue) (Fig. 3A). LX2 monocultures showed no detectable induction of α-SMA in response to 1PA2OA and ROL treatment (Fig. 3A, B). In contrast, LX2 cells co-cultured with Huh7 hepatocytes showed robust α-SMA expression, comparable to or exceeded the response induced by recombinant TGFb-1 (5ng/mL), the positive control for HSC activation (Fig. 3A, B). Strikingly, IsoA-transfected hepatocytes induced markedly higher α-SMA expression in LX2 cells compared with mock- or mHSD- transfected hepatocytes under both basal and supplemented conditions (Fig. 3A, B).

To further assess HSC activation, fibrogenic gene expression was quantified in LX2 cells isolated from co-cultures. *ACTA2* mRNA was strongly upregulated in LX2 cells co-cultured with IsoA-transfected hepatocytes under lipid-rich conditions, whereas mock- and mHSD-transfected hepatocytes failed to induce *ACTA2* in LX2 cells under the same conditions (Fig. 3C). Interestingly, *ACTA2* expression was comparable across co-cultures under basal conditions. In contrast, *COL1A1*, encoding the α1 chain of type I collagen, was selectively upregulated in LX2 cells co-cultured with IsoA-transfected hepatocytes under both basal and 1PA2OA plus ROL—supplemented conditions (Fig. 3D). Neither mock-nor mHSD-expressing hepatocytes induced *COL1A1* expression in either condition (Fig. 3D). Together, these findings reveal that HSD17B13 catalytic activity in hepatocytes is required for robust paracrine activation of HSCs in co-culture and induction of fibrogenic genes, delineating a functional hepatocyte– HSC axis that links hepatocellular lipid metabolism to fibrogenesis.

### HSD17B13-Dependent Secreted Factors from Hepatocytes Activate HSCs

To determine whether hepatocytes secrete soluble mediators that drive HSC activation, LX2 cells were treated with conditioned media (CM) collected from Huh7 cells transfected with mock, mHSD, or IsoA constructs. As controls, LX2 cells were cultured in basal medium, lipid-loaded medium, or treated with recombinant TGFb-1 (5ng/mL). Lipid supplementation alone did not induce LX2 activation, as evidenced by minimal changes in *ACTA2* (Fig. 4A) and *COL1A1* (Fig. 4B) expression relative to basal medium. In contrast, TGFb-1 robustly upregulated both *ACTA2* and *COL1A1* (Fig. 4A, B).

**Figure 4:**
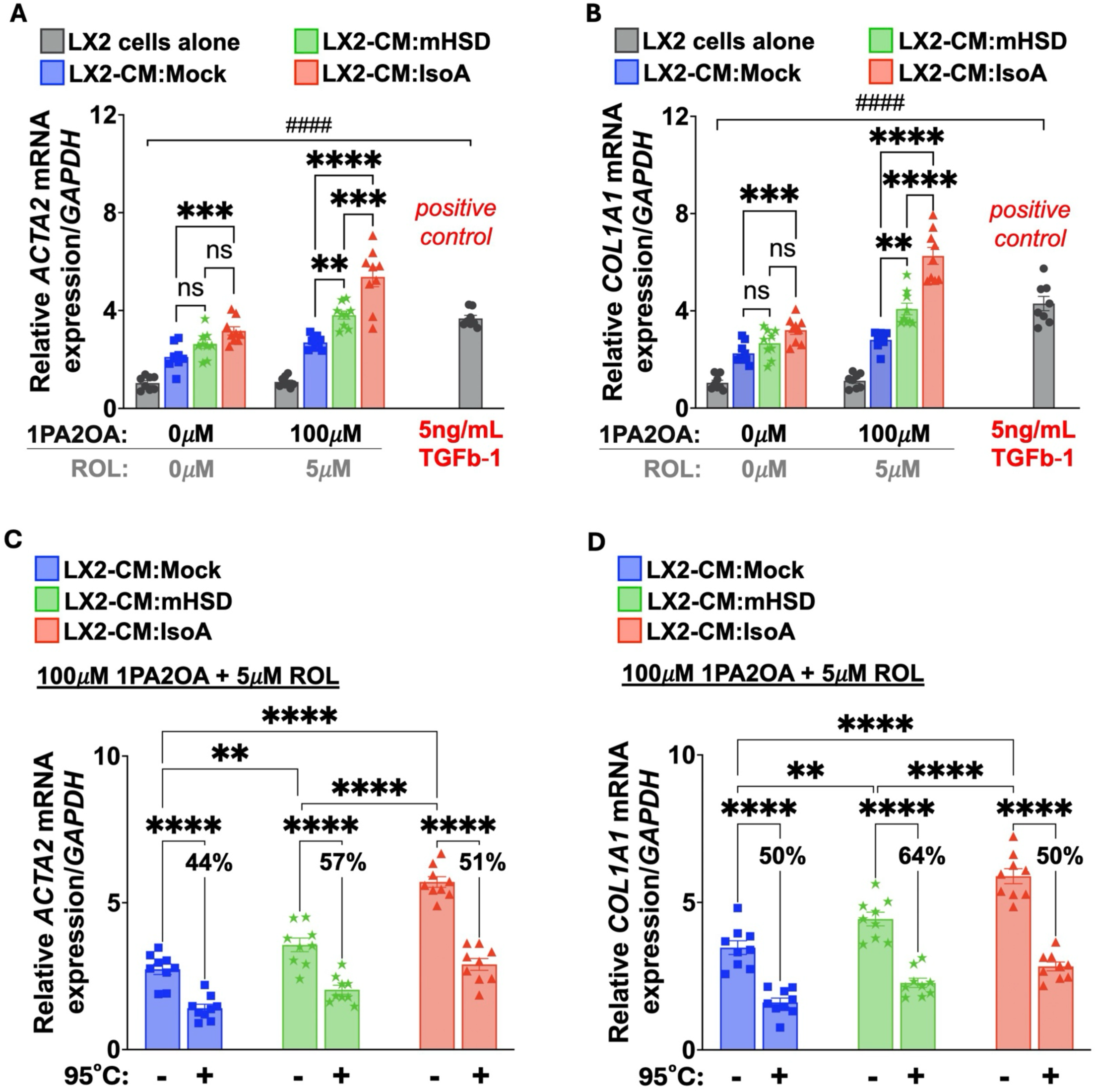
HSD17B13-dependent secreted factors from hepatocytes activate HSCs. **(A–B)** LX2 cells were treated with conditioned media (CM) collected from Huh7 hepatocytes transfected with mock, IsoA, or mHSD constructs to examine HSC activation by hepatocyte-derived secreted factors. LX2 cells cultured in basal medium, lipid-loaded medium or treated with recombinant TGFb-1 (5 ng/mL) served as controls. Lipid supplementation alone did not induce HSC activation, as evidenced by minimal *ACTA2* **(A)** and *COL1A1* **(B)** expression relative to basal medium. In contrast, TGFb-1 robustly upregulated both transcripts. Under lipid-loaded conditions, CM derived from IsoA- or mHSD-expressing hepatocytes significantly increased LX2 expression of *ACTA2* and *COL1A1* compared with mock-derived CM. Notably, mHSD-derived CM elicited a weaker induction of both genes relative to IsoA-derived CM. However, under basal conditions, CM from IsoA-but not mHSD-expressing hepatocytes induced LX2 expression of both markers compared with mock-derived CM. **(C–D)** Lipid-loaded CM from mock-, IsoA-, or mHSD-expressing hepatocytes was subjected to heat treatment (95°C for 20 min) prior to LX2 stimulation to evaluate whether the fibrogenic activity was heat-sensitive and protein-dependent. Heat treatment reduced the induction of *ACTA2* **(C)** and *COL1A1* **(D)** by ∼50% but did not abolish LX2 activation. Data are presented as mean ± SEM. (A—D) *n* = 9 biologically independent samples. Statistical significance was determined by one-way ANOVA followed by Tukey’s multiple-comparison test and by unpaired two-tailed Student’s *t*-test. ***ns*** *= P>0.05;* **\*** *= P≤0.05;* **\*\*** *= P≤0.01;* **\*\*\*** *= P≤0.001;* **\*\*\*\*** *= P≤0.0001*.

Under lipid-loaded conditions, CM derived from IsoA- or mHSD-expressing hepatocytes significantly increased LX2 expression of *ACTA2* and *COL1A1* (Fig. 4A, B) compared with mock-derived CM. By contrast, CM from IsoA- or mHSD-expressing hepatocytes cultured under basal conditions did not significantly upregulate these markers (Fig. 4A, B). Notably, mHSD-derived CM elicited a weaker induction of *ACTA2* and *COL1A1* relative to IsoA-derived CM (Fig. 4A, B). Together, these results indicate that HSD17B13 catalytic activity enhances the secretion of soluble profibrogenic mediators from hepatocytes, thereby promoting paracrine activation of HSCs under conditions of metabolic stress.

To test whether these mediators could be proteinaceous, lipid-loaded CM from mock-, mHSD-, or IsoA-expressing hepatocytes was heat-treated at 95 °C for 20 min prior to LX2 treatment. Heat treatment markedly attenuated the induction of *ACTA2* (Fig. 4C) and *COL1A1* (Fig. 4D) by approximately 50% but did not completely abolish LX2 activation. These findings suggest the possibility that the fibrogenic activity of hepatocyte-derived CM is mediated in part by heat-stable protein effectors, with potential contribution from non-protein factors. Altogether, these data demonstrate that HSD17B13 activity modulates the hepatocyte secretome to drive paracrine HSC activation under metabolic stress.

### HSD17B13 Activity Drives TGFb-1 Secretion from Lipid-Loaded Hepatocytes

To identify hepatocyte-derived factors potentially mediating HSD17B13-driven paracrine activation of HSCs, we profiled the expression of selected profibrotic and inflammatory genes in transfected Huh7 hepatocytes (Fig. 5A). Transfection efficiency was comparable between IsoA- and mHSD-transfected cells, with *HSD17B13* mRNA minimally expressed in mock controls (Fig. 5B). We next examined the mRNA expression of *TGFB1*, *TGFB2*, sonic hedgehog (*SHH*), hepatocyte nuclear factor 4α (*HNF4A*), interleukin-1β (*IL1B*), C-C motif chemokine ligand 5 (*CCL5*), and *CCL2* across group (Fig. 5A). *SHH*, *IL1B*, and *CCL2* expression remained unchanged across conditions, whereas *TGFB2*, *HNF4A*, and *CCL5* were modestly upregulated by both IsoA and mHSD expression. Strikingly, only *TGFB1* was significantly more elevated in IsoA-expressing cells compared with mHSD-expressing cells (Fig. 5A), suggesting that catalytically active HSD17B13 is necessary for optimal induction of *TGFB1* in hepatocytes, which parallels the enhanced HSC activation observed in IsoA co-cultures (Fig. 3A, B).

**Figure 5:**
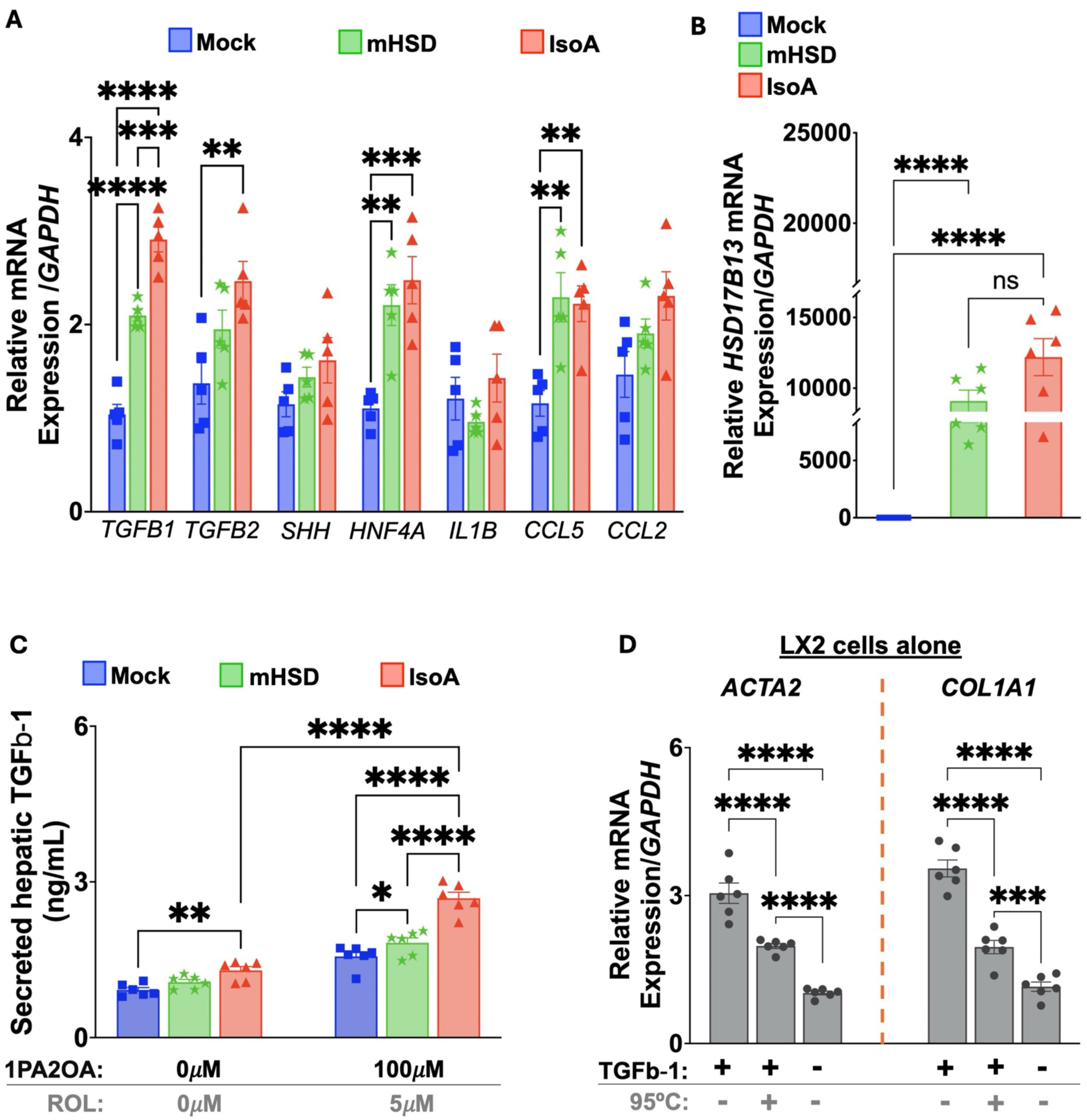
HSD17B13 activity drives TGFb-1 secretion from lipid-loaded hepatocytes. **(A)** Expression profiling of selected profibrotic and inflammatory genes in Huh7 hepatocytes transfected with mock, IsoA, or mHSD constructs revealed that among the genes examined (*TGFB1, TGFB2, SHH, HNF4A, IL1B, CCL5,* and *CCL2*), *TGFB1* mRNA was selectively and significantly upregulated in IsoA-expressing hepatocytes compared with mHSD, while other genes showed modest or no induction. **(B)** Quantitative RT-PCR confirmed comparable transfection efficiency between IsoA- and mHSD- expressing cells, with minimal *HSD17B13* expression in mock controls. **(C)** ELISA analysis demonstrated that IsoA overexpression markedly increased secretion of TGFb-1 protein into CM under lipid-loaded conditions (100 μM 1PA2OA + 5 μM ROL), mirroring the transcriptional upregulation of *TGFB1*. Lipid supplementation failed to induce substantial TGFb-1 secretion in mHSD-expressing hepatocytes. **(D)** Recombinant TGFb-1 (5 ng/mL) was heat-treated (95 °C for 20 min) prior to LX2 treatment to evaluate the stability of its fibrogenic activity. While untreated TGFb-1 robustly induced *ACTA2* and *COL1A1* expression in LX2 cells, heat treatment partially attenuated, but did not abolish these fibrogenic responses, shown by moderate decrease in *ACTA2* and *COL1A1*mRNA expression. Data are presented as mean ± SEM. (A—D) *n ≥* 5 biologically independent samples. Statistical significance was determined by one-way ANOVA followed by Tukey’s multiple-comparison post hoc test. ***ns*** *= P>0.05;* **\*** *= P≤0.05;* **\*\*** *= P≤0.01;* **\*\*\*** *= P≤0.001;* **\*\*\*\*** *= P≤0.0001*.

Based on these observations, we next evaluated whether catalytically active HSD17B13 promotes secretion of TGFb-1 protein into hepatocyte CM. ELISA analysis revealed that IsoA overexpression markedly increased TGFb-1 secretion, mirroring the transcriptional upregulation of *TGFB1* in hepatocytes (Fig. 5C). In contrast, lipid supplementation failed to induce substantial TGFb-1 secretion in mHSD-expressing cells (Fig. 5C), indicating that HSD17B13 catalytic activity is important for both transcriptional induction and secretion of TGFb-1 in metabolically stressed hepatocytes.

To assess the thermal stability of TGFb-1 and its fibrogenic activity, recombinant TGFb-1 (5 ng/mL) was heat-treated at 95 °C for 20 min prior to LX2 treatment. While untreated TGFb-1 robustly induced *ACTA2* and *COL1A1* expression, heat treatment attenuated, but did not abolish, fibrogenic responses in LX2 cells (Fig. 5D), consistent with the partial heat resistance observed in hepatocyte-derived CM (Fig. 4C, D). Together, these results identify TGFb-1 as a hepatocyte-derived paracrine signal regulated by HSD17B13 catalytic activity, that may mediate HSC activation during lipid overload.

### TGFb-1 Defines a Critical Axis of HSD17B13-Dependent Hepatocyte-HSC Crosstalk

To test whether TGFb-1 mediates HSD17B13-driven hepatocyte–HSC crosstalk (Fig. 3-5), we silenced *TGFB1* expression in hepatocytes using siRNA (siTGFb-1), as illustrated in the experimental schematic (Fig. 6A). Huh7 cells were transfected with mock, mHSD, or IsoA constructs, followed by siTGFb-1 treatment. Huh7 hepatocytes were then incubated with 100 μM 1PA2OA and 5 μM ROL for 48 h, and CM was collected. *TGFB1* expression was nearly abolished following siTGFb-1, achieving knockdown efficiencies of 86%, 90%, and 94% in mock-, mHSD-, and IsoA-expressing cells, respectively (Fig. 6B).

**Figure 6:**
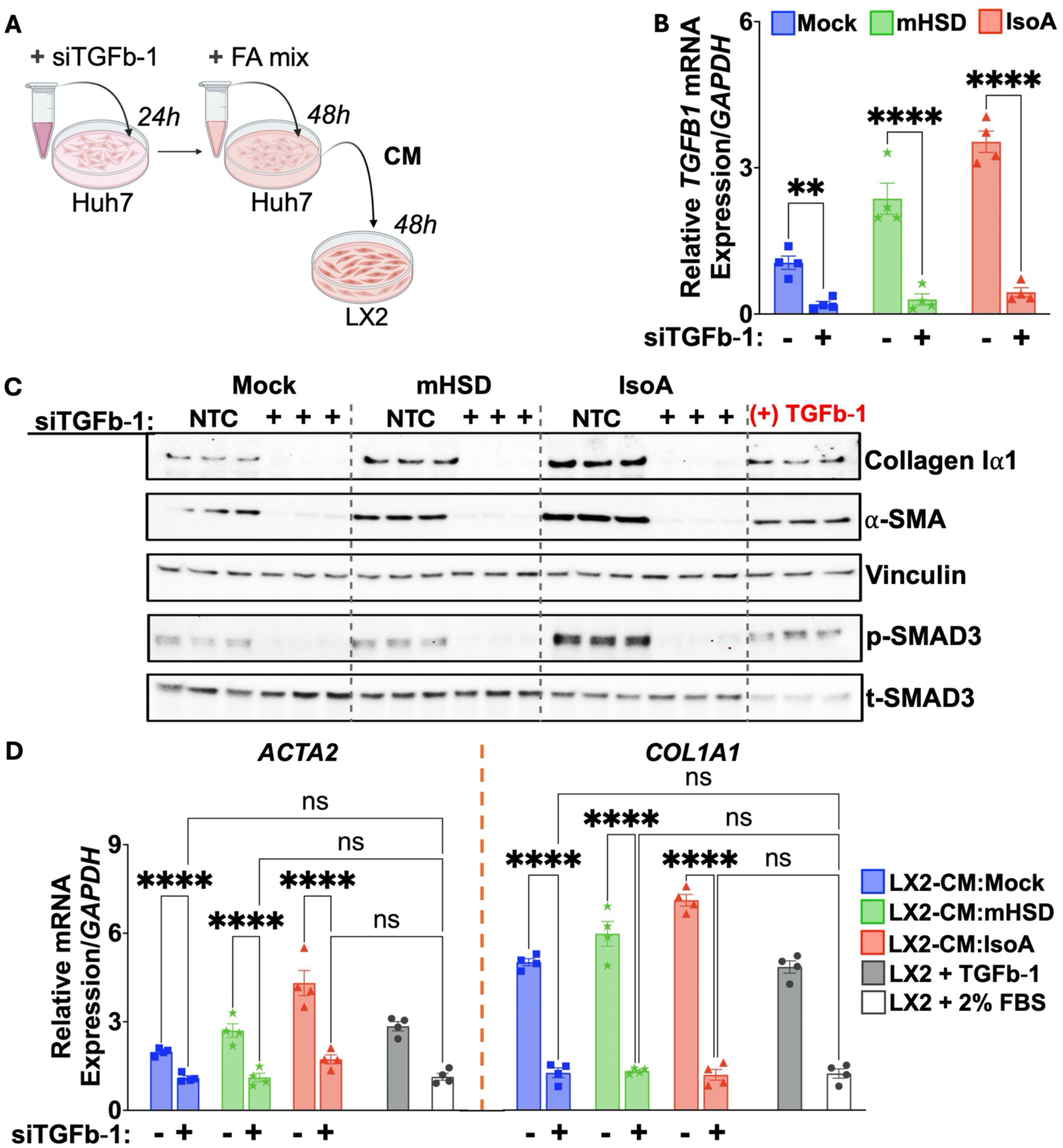
TGFb-1 defines a critical axis of *HSD17B13*-dependent hepatocyte–HSC crosstalk. **(A)** Experimental schematic illustrating the hepatocyte–HSC co-culture model incorporating *TGFB1* silencing. Huh7 hepatocytes were transfected with mock, IsoA, or mHSD constructs, followed by siRNA-mediated knockdown of *TGFB1* (siTGFb-1). Cells were subsequently cultured for 48 h in media containing 100 μM 1PA2OA and 5 μM ROL, after which CM were collected for LX2 stimulation. **(B)** Quantitative RT-PCR confirmed efficient *TGFB1* silencing, achieving knockdown efficiencies of 86%, 90%, and 94% in mock-, mHSD-, and IsoA-transfected hepatocytes, respectively. **(C)** Immunoblot analysis of LX2 cells treated with hepatocyte-derived CM revealed that IsoA-CM robustly induced collagen Iα1 and α-smooth muscle actin (α-SMA) protein expression, accompanied by increased phosphorylation of SMAD3 (pSMAD3). CM from mock- or mHSD- expressing hepatocytes elicited minimal induction. *TGFB1* knockdown in hepatocytes completely abolished the profibrotic effect of IsoA-CM by preventing the induction of collagen Iα1, α-SMA and the activation of SMAD3 signaling in LX2 cells. **(D)** Quantitative RT-PCR analysis of *ACTA2* and *COL1A1* expression in LX2 cells confirmed that *TGFB1* silencing normalized fibrogenic gene expression to baseline levels observed in quiescent HSCs. Data are presented as mean ± SEM. (B—E) *n* ≥ 4 biologically independent samples. The unpaired two-tailed Student’s *t*-test was performed to establish statistical significance between groups. ***ns*** *= P>0.05;* **\*** *= P≤0.05;* **\*\*** *= P≤0.01;* **\*\*\*** *= P≤0.001;* **\*\*\*\*** *= P≤0.0001*.

**Figure 7:**
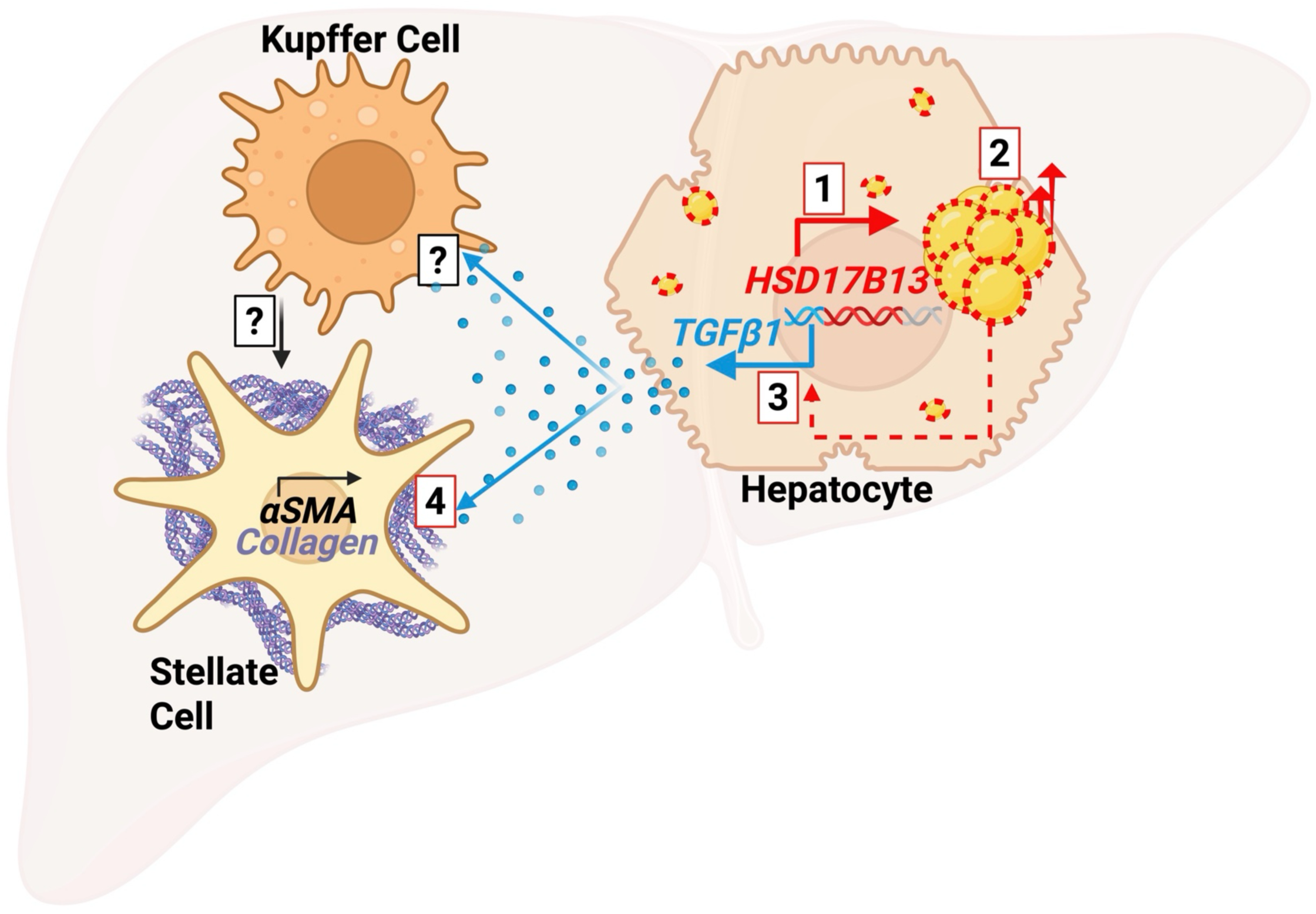
HSD17B13-mediated HSC activation *in vitro* is hepatocyte TGFb-1-dependent. Schematic model summarizing the proposed mechanism by which *HSD17B13* promotes LD remodeling and TGFb-1 secretion, thereby coupling hepatocellular lipid metabolism and metabolic stress to paracrine fibrogenic signaling. **1)** Hepatocyte *HSD17B13* expression is upregulated under lipid-loading conditions. **2)** HSD17B13 localizes to the LD surface, where HSD17B13 promotes TG storage and hepatocellular lipid accumulation through its enzymatic and potential scaffolding functions. **3)** Catalytically active HSD17B13 stimulates hepatocyte *TGFB1* transcription and TGFb-1 protein secretion, potentially involving pathways such as platelet-activating factor (PAF) biosynthesis or other lipid-derived intermediates. **4)** Secreted hepatocyte TGFb-1 acts on neighboring HSCs to activate canonical TGFb-1/SMAD3 signaling, inducing α-SMA and collagen Iα1 expression, encoded by *ACTA2* and *COL1A1*, respectively. Hepatocyte-derived TGFb-1 may also recruit inflammatory cells and perpetuate chronic inflammation, ultimately reinforcing HSC activation and ECM deposition under conditions of lipid overload and metabolic stress. This model is based on statistically validated findings described in Figures 1–6. Statistical significance was determined by one-way ANOVA with Tukey’s post hoc tests or unpaired two-tailed Student’s *t*-test. ***ns*** *= P>0.05;* **\*** *= P≤0.05;* **\*\*** *= P≤0.01;* **\*\*\*** *= P≤0.001;* **\*\*\*\*** *= P≤0.0001*.

Consistent with earlier findings, CM from IsoA-transfected hepatocytes induced robust protein expression of collagen Iα1 (encoded by *COL1A1*) and α-SMA in LX2 cells, accompanied by marked phosphorylation of SMAD3 (pSMAD3) (Fig. 6C; Fig. S4A, B). By contrast, CM from mock- or mHSD- expressing cells elicited minimal response in LX2 cells (Fig. 6C). Furthermore, pSMAD3 levels in LX2 cells exposed to mock- or mHSD-derived CM were comparable (Fig. S4B), suggesting that HSD17B13 catalytic activity likely stimulates fibrogenic response in HSCs by driving hepatocyte TGFb-1 secretion and subsequent activation of canonical SMAD3 signaling in HSCs.

Importantly, silencing *TGFB1* in hepatocytes completely abolished the profibrotic effect of catalytically active HSD17B13 and prevented downstream activation of HSCs, as evidenced by loss of collagen Iα1, α-SMA, and pSMAD3 induction in LX2 cells (Fig. 6C; Fig. S4A, S4B). Furthermore, *TGFB1* knockdown suppressed HSD17B13-induced upregulation of *ACTA2* and *COL1A1* transcripts in LX2 cells, reversing expression to baseline levels observed in quiescent HSCs (Fig. 6D). These results establish TGFb-1 as a critical paracrine mediator of HSD17B13-driven HSC activation in this model system.

To further validate this finding, TGFb-1 in hepatocyte CM was neutralized with a blocking antibody (α- TGFb-1). CM from IsoA-expressing hepatocytes strongly induced fibrotic genes in LX2 cells, whereas mock- and mHSD-derived CM failed to elicit a robust fibrogenic response in LX2 cells. Notably, TGFb- 1 neutralization markedly reduced IsoA-CM-induced expression of collagen Iα1 and α-SMA (Fig. S4C, S4D). Collectively, these results demonstrate that the enzymatic function of HSD17B13 is required for enhanced TGFb-1 secretion from lipid-laden hepatocytes, which in turn activates HSCs via canonical TGFb-1/SMAD3 signaling. Remarkably, the complete loss of fibrogenic activity in LX2 cells following TGFb-1 inhibition indicates that HSD17B13-driven fibrogenesis is mediated primarily by TGFb-1 in our experimental system, either as the dominant driver of HSC activation or as a permissive signal that primes HSCs to respond to other hepatocyte-derived mediators.

## Discussion

Major interest in *HSD17B13* as a regulator of human MASH stems from multiple genetic studies consistently associating *HSD17B13* with liver disease progression, showing naturally occurring LoF variants conferring protection against steatohepatitis and fibrosis across diverse populations [36–43]. Recent work challenges the interpretation of these alleles as complete LoF, as some truncated variants retain partial catalytic activity [79]. Nevertheless, the well-characterized splice variant *rs72613567* yields a truncated HSD17B13 protein that is destabilized and rapidly degraded, resulting in reduced abundance and functional capacity [36]. Thus, despite residual activity, the net reduction in bioavailable HSD17B13 likely limits its pathogenic capacity. Notably, *HSD17B13* expression is largely hepatocyte-specific and is further induced in MASH [36, 46, 60, 80]. Yet the clinical phenotype most strongly associated with *HSD17B13* LoF is reduced liver fibrosis, a major determinant of adverse outcomes in MASH and a process driven primarily by HSCs [36, 81]. These observations underscore a central gap in our understanding of how hepatocyte-restricted HSD17B13 influences HSC activation and fibrogenic signaling.

We identify hepatocyte-derived TGFb-1 as a principal mediator linking *HSD17B13* expression to HSC activation *in vitro*. (Figs. 5 and 6). By fine-tuning IsoA transcript expression to physiologic levels observed in PHHs (Fig. 1A), we show that HSD17B13 robustly induces *TGFB1* transcription and secretion (Fig. 5). TGFb-1 is recognized as a master profibrogenic cytokine that initiates HSC activation and sustains their myofibroblast-like phenotype, driving ECM deposition and tissue remodeling [71, 72, 74, 82–85]. Elevated hepatic or circulating TGFb-1 levels correlate with fibrosis severity, progression to cirrhosis, and adverse clinical outcomes [73, 86, 87]. Given these observations and the potent LX2 activation induced by IsoA-CM (Fig. 4), we determined whether hepatocyte-derived TGFb-1 mediates the profibrogenic effects of HSD17B13 in co-culture. Both *TGFB1* silencing (Fig. 6) and antibody-based neutralization of TGFb-1 (Fig. S4) completely abolished LX2 activation. The abrogation of LX2 activation upon TGFb-1 depletion is striking given the complexity of the hepatocyte secretome. Our finding rules out the possibility that TGFb-1 merely amplifies other mediators (Fig. S4) and instead indicates that TGFb-1 is a necessary node in HSD17B13-driven paracrine LX2 activation in our system. Together, these results highlight steatotic hepatocytes as active contributors to the fibrotic niche and identify TGFb-1 as a dominant profibrogenic output of HSD17B13 in this model.

The *in vivo* fibrotic milieu is considerably more complex. Fibrogenesis arises from coordinated interactions among hepatocytes, HSCs, and immune cells that collectively sustain chronic inflammation and ECM accumulation. Hepatocyte-derived TGFb-1 and other stress signals (e.g., DAMPs, mitochondrial DNA, and oxidized lipids) can activate Kupffer cells and infiltrating macrophages, which in turn amplify fibrogenic responses by secreting additional inflammatory and profibrogenic mediators [85, 88–95]. Beyond intrahepatic sources, platelet-derived TGFb-1 also promotes HSC activation and collagen deposition, whereas platelet depletion or genetic inactivation of platelet TGFb-1 protects against experimental hepatic fibrosis [96]. Together, this evidence highlights TGFb-1 signaling as a central driver of HSC activation and progressive fibrosis. TGFb-1 is secreted as a latent complex bound to latency-associated peptide (LAP) and ECM proteins. Upon activation, TGFb-1 binds TGFb receptor type II (TGFbRII) to initiate SMAD-dependent transcription of collagen and other ECM-related genes [70, 72, 73, 75, 76, 85]. Importantly, HSC-specific conditional deletion of *Tgfbr2* markedly reduces fibrosis and improves metabolic and inflammatory profiles in obese mice [97], underscoring the critical role of HSC-specific TGFb-1/TGFbRII signaling in MASH-associated fibrogenesis. However, while TGFb-1 remains an appealing antifibrotic target, its systemic inhibition is challenging given its critical roles in immune regulation and tumor suppression [96, 98].

Our results also provide insights into how HSD17B13 regulates lipid turnover. Although HSD17B13 has reported activity towards hydroxysteroids, eicosanoids, and retinoids [36, 62], its contribution to lipid handling and metabolic regulation has remained unclear due to conflicting reports. Ma et al. (2019) observed no effect of *HSD17B13* knockout or overexpression on lipid content in FA-loaded hepatocytes, whereas other studies reported increased LD accumulation with HSD17B13 overexpression [45, 68, 99]. Our data show that overexpression of enzymatically active HSD17B13 (IsoA) robustly enhanced neutral lipid accumulation (Fig. 1, S1), whereas a catalytically inactive mutant (S172A, mHSD), despite localizing to LDs, failed to promote steatosis (Fig 2). Together, these findings indicate that HSD17B13 catalytic function is required to drive hepatocellular lipid accumulation, which likely contributes to *TGFB1* upregulation.

Mechanistically, HSD17B13 expression was associated with reduced glycerol release from hepatocytes (Fig. S3), consistent with prior reports [68]. However, this reduction was also observed with mHSD expression, indicating that suppression of lipolysis alone does not account for the increased intracellular lipids by catalytically active HSD17B13 (Figs. 1, 2). However, IsoA expression markedly upregulated ChREBP and induced several DNL-related enzymes (Fig. S3), whereas mHSD was less effective at inducing these markers. These results suggest that HSD17B13 catalytic activity is required to achieve maximal lipid accumulation, at least in part, by promoting lipogenic transcriptional programs. IsoA expression also increased full-length SREBP-1 but did not affect its proteolytic maturation, in contrast to murine studies that link Hsd17b13 to SREBP-1 processing [69], highlighting potential species-specific differences that warrant careful consideration when translating findings to human disease.

Our findings raise important mechanistic questions about how HSD17B13 couples LD accumulation to TGFb-1 production. An intriguing avenue for future research involves liquid–liquid phase separation (LLPS) of HSD17B13 on LD surfaces. HSD17B13 has been shown to form LLPS condensates through its N-terminal intrinsically disordered region, which enables dynamic self-clustering in hepatocytes and correlates with enhanced enzymatic activity [100]. Such condensates have also been observed in livers of patients with MASH [100], supporting a potential pathophysiological role for this LLPS-dependent organization. We propose that phase-separated HSD17B13 condensates act as localized enzymatic hubs that accelerate phospholipid turnover, particularly platelet-activating factor (PAF) biosynthesis and stimulate arachidonic acid (AA) release [100, 101]. Although hepatocytes lack canonical PAF receptors, PAF can signal through receptor-independent pathways to mobilize AA, which is subsequently metabolized into eicosanoids and reactive oxygen species (ROS) [101–106]. These ROS species can promote *TGFB1* transcription through redox-sensitive transcriptional pathways and facilitate post-translational activation of latent TGFb-1, thereby expanding the pool of bioactive TGFb-1 [107–113]. Beyond lipid mediator pathways, retinoic acid (RA) metabolism may provide an additional regulatory layer linking HSDB17B13 to TGFb-1 synthesis. Catalytically active HSD17B13 catalyzes the oxidation of retinol to retinaldehyde, facilitating RA production via ALDH1A1, which is upregulated in our system [62]. RA signaling through RA-responsive nuclear receptors (RAR/RXR) can enhance *TGFB1* transcription [114], suggesting that HSD17B13-dependent RA production may establish an autocrine feedback loop that potentially reinforces *TGFB1* expression under lipid stress. If validated, this model would provide a mechanistic framework linking HSD17B13 enzymatic activity to transcriptional programs that drive fibrogenesis. Future studies should directly compare the effect of wild-type and catalytically inactive HSD17B13 on condensate formation, RA signaling, PAF synthesis, AA release, ROS generation, and TGFb-1 activation, and incorporate pharmacologic inhibition of AA metabolism and RA signaling to delineate their respective contributions.

In summary, our study expands the paradigm of LD biology by redefining HSD17B13 as a catalytic hub that links LD expansion to TGFb-1 production and HSC activation. This work establishes a conceptual framework for understanding how hepatocyte-restricted HSD17B13 drives fibrogenic signaling and contributes to MASH pathogenesis.

## Acknowledgements

We are grateful to the members of the Czech laboratory for their valuable discussions and to the UMASS Chan Metabolic Disease Research Center for expert support with methodologies and metabolic profiling. We also thank Kerri Miller and Sarah Nicoloro for administrative assistance. This work was supported by the National Institutes of Health (National Institute of Diabetes and Digestive and Kidney Diseases (NIDDK) grants DK103047 and DK130852).

## Author Contributions

N.R., B.Y., and M.P.C. conceived the study and developed the experimental design and methodology. N.R. performed experiments, analyzed data, and prepared figures. L.L. quantified lipid droplet volumes from confocal images using ImageJ. K.M. contributed to methodology development. C.C. and G.I. performed mass spectrometry-based quantification and analysis of retinoid species. N.R. and M.P.C. wrote the manuscript, with input from B.Y. All authors reviewed and approved the final version of the manuscript.

## Supplemental Data

**Table 1:**
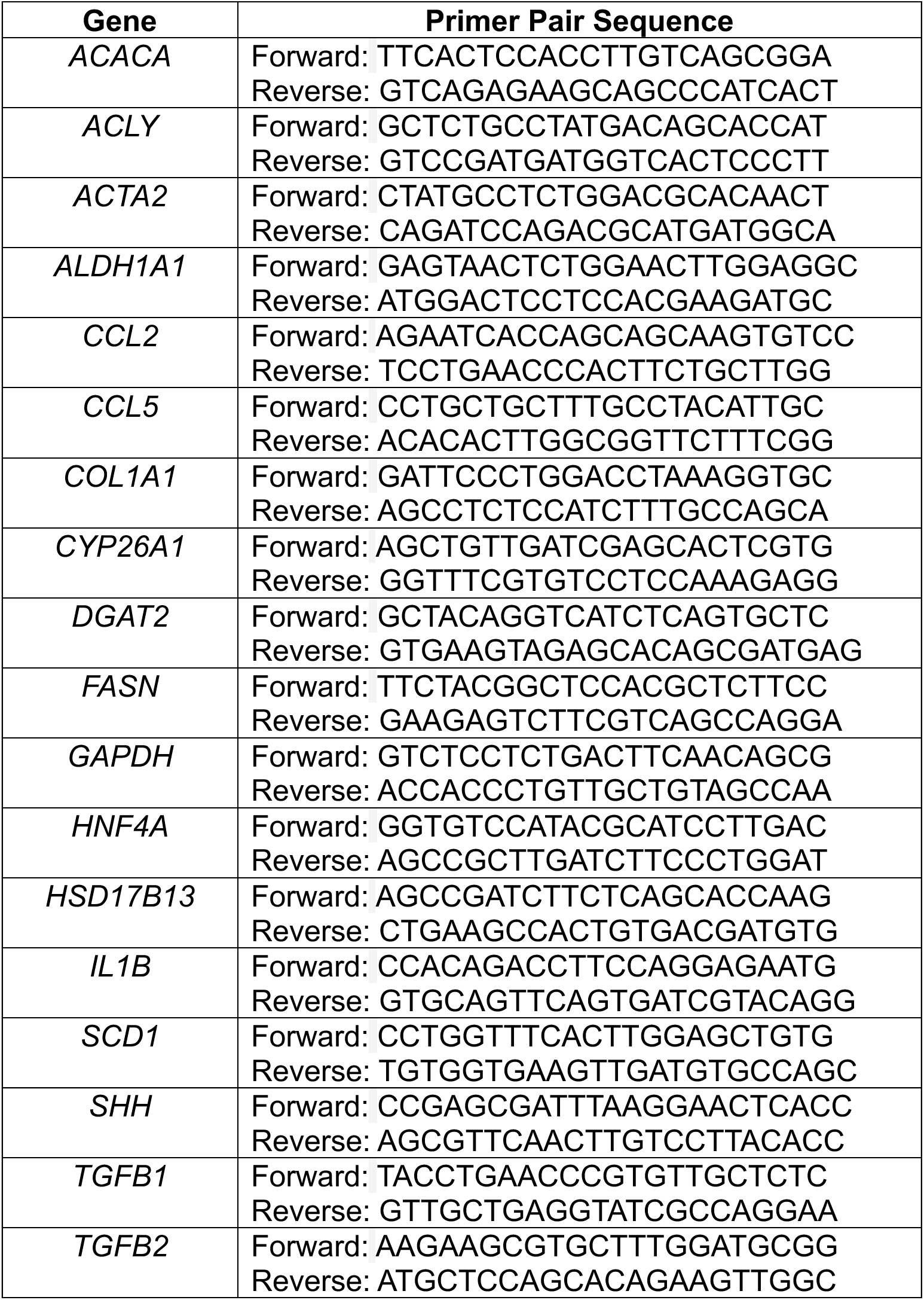
Human primers used for RT-qPCR.

**Table 2:** Immunoblotting reagents used in this study.

| **Blotting Probe** | **Supplier** | **Cat. #** |
| --- | --- | --- |
| Anti-α-SMA | Abcam | Ab124964 |
| Anti-ChREBP | Novus Biological | NB400-135 |
| Anti-Collagen type I | Southern Biotech | 1310-01 |
| Anti-HA-Tag-HRP | Thermo Fisher | PA1-29751 |
| Anti-Histone H3 | Cell Signaling Technology | 4499S |
| Anti-HSD17B13 | Thermo Fisher | PA5-25633 |
| Anti-HSD17B13 (IF) | Thermo Fisher | PA5-56063 |
| Anti-HSP90-HRP | Cell Signaling Technology | 79641S |
| Anti-Phospho-SMAD3 | Cell Signaling Technology | 9520S |
| Anti-SMAD3 | Cell Signaling Technology | 9523S |
| Anti-SREBF1 | Protein Tech | 14088-1-AP |
| Anti-Vinculin | Cell Signaling Technology | 18799 |
| Goat Anti-Rabbit-647 | Thermo Fisher | A-21244 |
| Goat Anti-Rabbit IgG-HRP | Invitrogen | 65-6120 |
| Mouse Anti-Goat IgG-HRP | Santa Cruz | sc-2354 |
| Mouse IgG1 isotope control Ab | Thermo Fisher | 02-6100 |
| TGFb-1,2,3 monoclonal Ab (1 μg/mL) | Thermo Fisher | MA5-23795 |

**Supplemental Figure 1:**
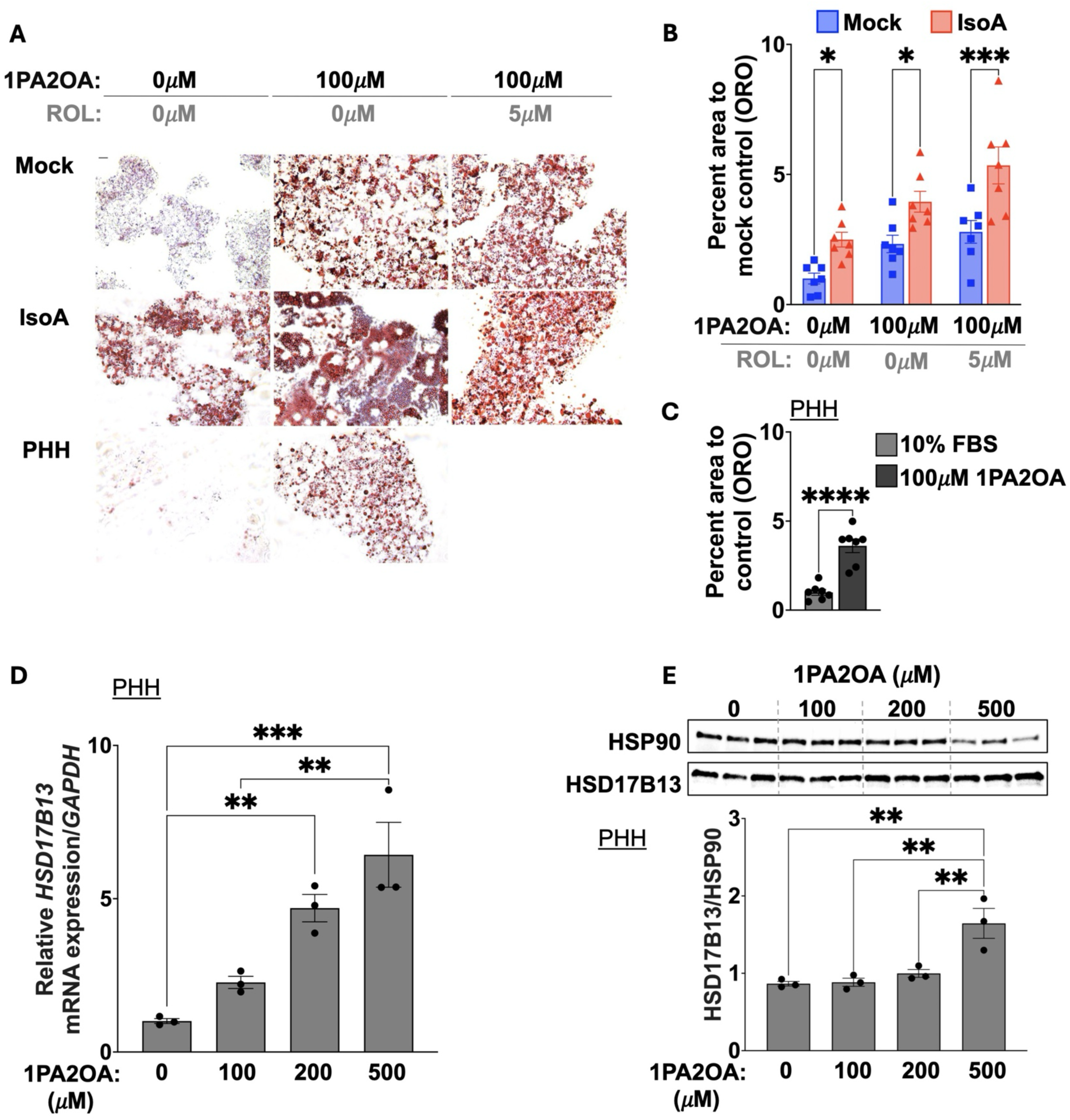
HSD17B1*3* promotes lipid accumulation across human hepatoma and primary hepatocyte systems. **(A–B)** HepG2 cells transfected with mock or IsoA constructs, and PHH were cultured under basal or lipid-loading conditions as indicated. Cells were fixed and stained with Oil Red O (ORO) to visualize neutral lipids (scale bar, 50μm). IsoA overexpression in HepG2 cells markedly increased intracellular lipid accumulation under both basal and lipid-supplemented conditions. **(C)** In PHHs, lipid accumulation was minimal under basal culture conditions but increased significantly following 100 μM 1PA2OA treatment. **(D–E)** Dose–response experiments in PHHs demonstrated that increasing 1PA2OA concentrations progressively enhanced lipid accumulation and induced *HSD17B13* mRNA expression, whereas HSD17B13 protein levels remained largely unchanged at concentrations below 500 μM. Data are presented as mean ± SEM. (D—E) *n* = 3 biologically independent samples. Statistical significance between groups was determined by ordinary one-way ANOVA followed by Turkey’s multiple-comparison post hoc test and by unpaired two-tailed *t*- test. ***ns*** *= P>0.05;* **\*** *= P≤0.05;* **\*\*** *= P≤0.01;* **\*\*\*** *= P≤0.001;* **\*\*\*\*** *= P≤0.0001*.

**Supplemental Figure 2:**
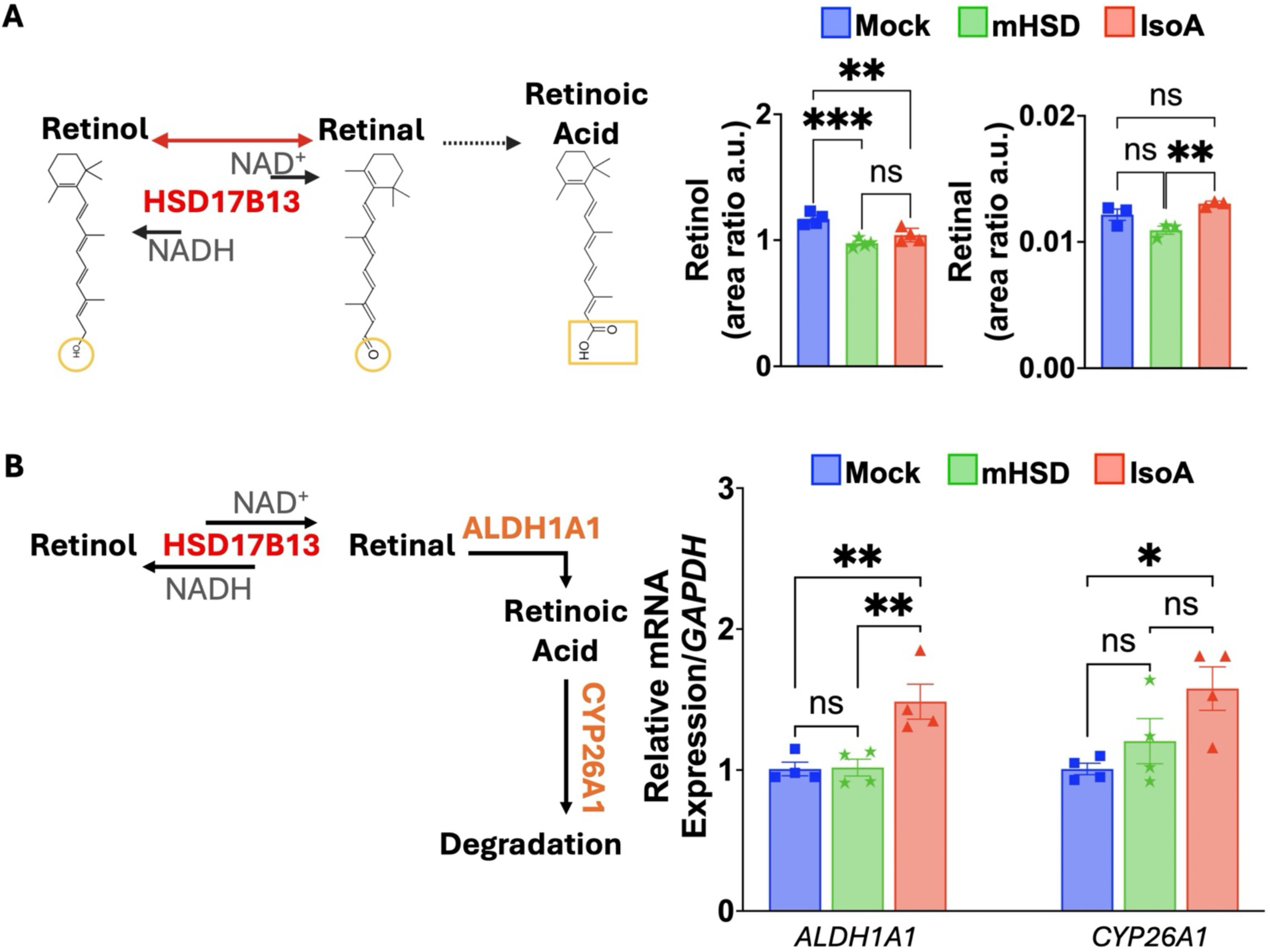
The S172A mutation impaired the catalytic activity of HSD17B13. **(A)** Schematic representation of retinol metabolism to retinoic acid (RA), highlighting the oxidation of retinol to retinaldehyde catalyzed by HSD17B13. Mass spectrometry analysis and quantification of extracted retinoids (retinol and retinaldehyde) from transfected hepatocytes revealed that the catalytically inactive mutant (mHSD) exhibited markedly reduced enzymatic activity toward retinol compared with IsoA in a cell-based retinol conversion assay. **(B)** Expanded pathway schematic illustrating the coordinated function of HSD17B13, ALDH1A1, and CYP26A1 in retinoid metabolism. Enzyme positions are labeled along the conversion steps, with HSD17B13 between retinol and retinal, ALDH1A1 between retinal and RA, and CYP26A1-mediated RA degradation. Quantitative RT-PCR showed that IsoA overexpression upregulated mRNA expression of *ALDH1A1* and *CYP26A1* in Huh7 cells, consistent with activation of RA-responsive feedback regulation of retinol metabolism. Data presented as mean ± SEM. (A—B) *n* ≥ 3 biologically independent samples. Statistical significance was determined by ordinary one-way ANOVA followed by Tukey’s multiple-comparison post hoc test. ***ns*** *= P>0.05;* **\*** *= P≤0.05;* **\*\*** *= P≤0.01;* **\*\*\*** *= P≤0.001;* **\*\*\*\*** *= P≤0.0001*.

**Supplemental Figure 3:**
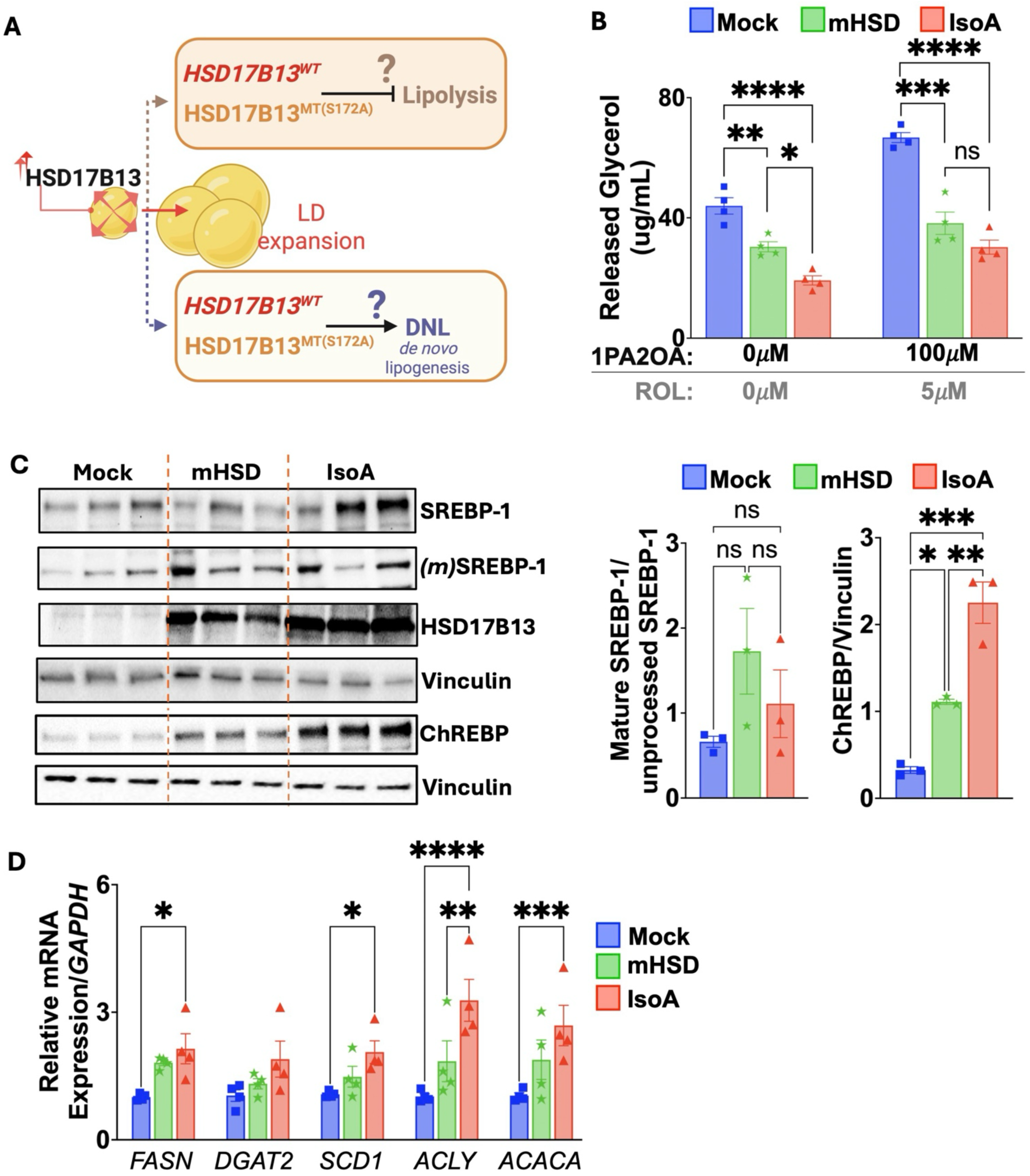
HSD17B13-driven activation of lipogenic pathways requires catalytic activity. **(A)** Graphical Schematic overview of LD turnover highlighting the likely contribution of HSD17B13 to the imbalance between lipid synthesis (de novo lipogenesis, DNL) and lipolysis to promote LD expansion. **(B)** Lipolytic activity was assessed by quantifying glycerol release in transfected Huh7 cells. Under lipid-loading conditions, glycerol release was significantly reduced in both IsoA- and mHSD-expressing cells. Under basal conditions, however, IsoA expression led to a more pronounced impairment of lipolytic activity, indicating that HSD17B13, in part, suppresses lipolysis independently of catalytic activity. **(C)** Immunoblot analysis of sterol regulatory element-binding protein-1 (SREBP-1) and carbohydrate response element-binding protein (ChREBP) was performed to evaluate DNL activation. No significant changes were observed in the ratio of mature (mSREBP-1) to full-length SREBP-1 among groups, whereas ChREBP protein abundance was markedly increased in IsoA-transfected cells compared with mock and mHSD. **(D)** Quantitative RT-PCR analysis of DNL-associated genes showed that IsoA, but not mHSD, significantly upregulated *FASN*, *SCD1*, *ACACA*, and *ACLY* expression, consistent with enhanced lipogenic signaling. Data are presented as mean ± SEM. (B—D) *n* ≥ 3. Statistical significance was determined by one-way ANOVA followed by Tukey’s multiple-comparison post hoc test. ***ns*** *= P>0.05;* **\*** *= P≤0.05;* **\*\*** *= P≤0.01;* **\*\*\*** *= P≤0.001;* **\*\*\*\*** *= P≤0.0001*.

**Supplemental Figure S4:**
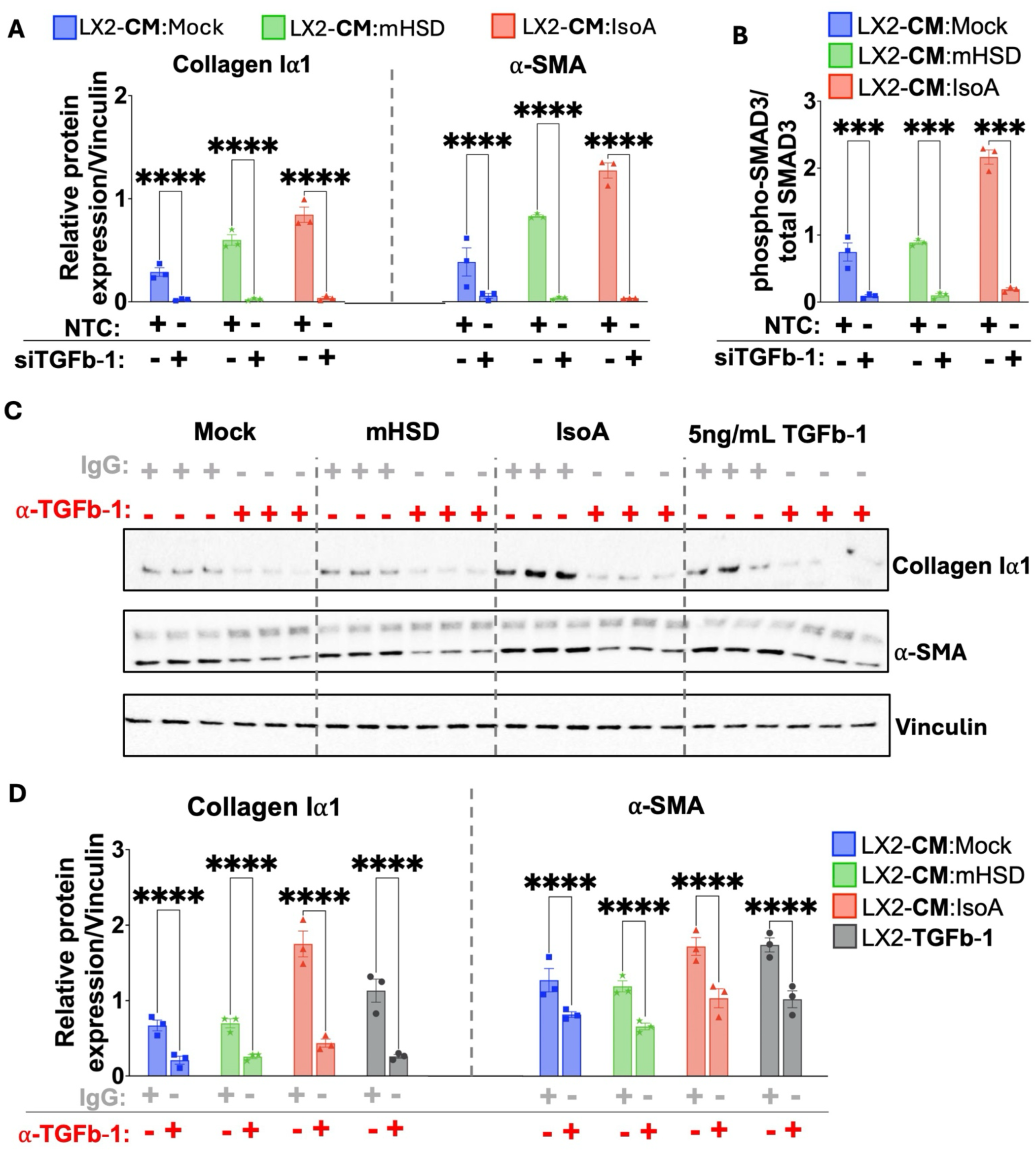
HSD17B13-dependent activation of HSCs occurs through TGFb- 1/SMAD3 signaling and is abolished by TGFb-1 neutralization. **(A–B)** Immunoblot validation of LX2 responses showing that IsoA-derived CM induced strong α-SMA, collagen Iα1, and pSMAD3 expression compared with CM from mock- or mHSD-transfected hepatocytes. Following *TGFB1* silencing, induction of these markers in LX2 cells was completely abolished. **(C–D)** Neutralization of TGFb-1 in IsoA-derived CM using a blocking antibody (α-TGFb-1; 1 μg/mL) markedly reduced IsoA-CM–induced expression of collagen Iα1 and α-SMA in LX2 cells, phenocopying the effects of *TGFB1* knockdown. Data are presented as mean ± SEM. (A—D) *n* = 3 biologically independent samples. The unpaired two-tailed Student *t*-test was performed to establish statistical significance between groups. ***ns*** *= P>0.05;* **\*** *= P≤0.05;* **\*\*** *= P≤0.01;* **\*\*\*** *= P≤0.001;* **\*\*\*\*** *= P≤0.0001*.

